# Cell-preferential PGRL1 paralogs provide distinct modes of photoprotection in C_4_ photosynthesis

**DOI:** 10.1101/2024.11.28.625945

**Authors:** Samuel J. Nix, Craig J. Morton, Russell Woodford, Robert T. Furbank, Maria Ermakova

## Abstract

The PGR5-PGRL1 pathway protects plants from photodamage by regulating electron flow to maintain Photosystem I in an oxidised state. Grasses possess two PGRL1 paralogs, but their functional roles have remained unknown. Here, we show that the ancestral PGRL1 paralog, PGRL1α, which is conserved across algae and land plants, is enriched in the mesophyll cells of grasses that perform the NADP-ME subtype of C_4_ photosynthesis. In contrast, the grass-specific paralog PGRL1β is enriched in bundle sheath cells. To investigate the functional significance of this cell-specific expression, we generated gene-edited lines of the NADP-ME C_4_ grass *Setaria viridis* lacking either PGRL1 paralog. We found that PGRL1β in bundle sheath cells was required for rapid photoprotection, enabling Photosystem I oxidation five seconds faster during transitions from darkness to high light. In contrast, PGRL1α in mesophyll cells was essential for maintaining Photosystem I oxidation under steady-state high-light conditions. We propose that these complementary functions arise from structural differences within the lumen-facing regions of the two paralogs, providing a mechanistic basis for their distinct roles in regulating photoprotection. The conservation of this dual PGRL1 system across grasses suggests that functional specialisation of the paralogs expands the dynamic range of protective responses, enhancing photosynthetic performance under fluctuating and high-light stress conditions.

## Introduction

Light is essential for photosynthesis, and therefore for plant productivity, but an excess of absorbed light can over-reduce the chloroplast electron transport chain and damage the photosynthetic machinery. Plants have evolved multiple strategies, collectively known as photoprotection, that prevent over-reduction of the chloroplast electron transport chain and help avoid damage of Photosystem I (PSI), which, unlike damage of Photosystem II (PSII), is irreversible (Tikkanen & Aro, 2014; Walker *et al*., 2020). Trans- thylakoid proton motive force (*pmf*), or more specifically the proton gradient (ΔpH) component of *pmf,* plays a critical role in regulating photoprotection. *Pmf* is established largely by the Cytochrome *b*_6_*f* complex (Cyt*b*_6_*f*) which couples electron transport with shuttling protons from the chloroplast stroma to the thylakoid lumen. While *pmf* is essential for powering ATP synthase activity to generate ATP, when proton influx into the lumen exceeds efflux, lumen acidification below pH 6 triggers two key mechanisms of photoprotection: energy-dependent non-photochemical quenching (qE) and photosynthetic control (Kramer *et al*., 1999). qE dissipates excess absorbed light energy as heat in the Photosystem II (PSII) antennae, while photosynthetic control slows down Cyt*b*_6_*f* activity to restrict electron flux to PSI (Li *et al*., 2004; Degen & Johnson, 2024).

PROTON GRADIENT REGULATION 5 (PGR5) and PROTON GRADIENT REGULATION 5-LIKE 1 (PGRL1) form a pathway essential for pH-dependent photoprotection. Plants lacking either PGR5 or PGRL1 suffer severe PSI damage under fluctuating light due to a diminished ability to establish *pmf*, qE and photosynthetic control (Munekage *et al*., 2002; Suorsa *et al*., 2012; Yamori *et al*., 2016; Woodford *et al*., 2025). Molecular studies suggest that transmembrane PGRL1 and soluble PGR5 form a heterodimer complex (DalCorso *et al*., 2008; Hertle *et al*., 2013). PGR5 appears to regulate electron transport while PGRL1 mainly acts as a gatekeeper of PGR5 function and prevents PGR5’s degradation by PGRL2 (Rühle *et al*., 2021; Chaturvedi *et al*., 2024). In the dark, PGRL1 is dimerised via a disulphide bond between conserved stromal cysteines which keeps PGR5 inactive, while the reduction of PGRL1’s disulphide bond by Thioredoxin *m*4 (Trxm4) under light activates PGR5 (Hertle *et al*., 2013; Okegawa & Motohashi, 2020; Chaturvedi *et al*., 2024). Although mechanistic details remain unresolved, the PGR5-PGRL1 pathway is proposed to build up *pmf* by facilitating cyclic electron flow (CEF) around PSI, i.e. directing electrons from Ferredoxin back to Cyt*b*_6_*f* (Munekage *et al*., 2002; Joliot & Johnson, 2011).

In contrast to most dicots where PGRL1 is encoded by a single gene, some grasses are reported to contain two *PGRL1* homologs (Zhang *et al*., 2024). It is also noted that transcripts of one of these homologs appear to be more abundant in bundle sheath (BS) cells compared to mesophyll (M) cells in *Zea mays* (maize), *Sorghum bicolor* (sorghum), and *Setaria viridis* – grasses that employ NADP-ME subtype of C_4_ photosynthesis (Wu & Guo, 2023). C_4_ photosynthesis provides a strong advantage over conventional C_3_ photosynthesis in low CO₂, high temperature and dry environments (Sage *et al*., 2011). This is achieved through operation of the C_4_ metabolic cycle across M and BS cells which increases CO_2_ partial pressure in BS cells where the main CO_2_ fixing enzyme, ribulose-1,5-bisphosphate carboxylase/oxygenase (Rubisco), resides (Hatch, 1987; von Caemmerer & Furbank, 2003). Most agronomically critical C_4_ grasses belong to the NADP-ME subtype, predominantly relying on NADP^+^-dependent malic enzyme for decarboxylation, but two other subtypes, NAD-ME and PCK, as well as combinations of subtypes also exist (Furbank, 2011; Ermakova *et al*., 2026). A distinct feature of NADP-ME plants is that decarboxylation of malate, produced in M cells with the use of NADPH, generates NADPH in BS chloroplasts. This means that while M chloroplasts require both PSII and PSI, BS chloroplasts have lower PSII activity and mostly rely on PSI (Chapman *et al*., 1980; Ermakova *et al*., 2021c).

Here we integrated phylogenetics, structural modelling, cell-specific expression data, and physiological analysis of gene-edited *Setaria viridis* to identify the roles of the two PGRL1 paralogs. We demonstrate that while functions of PGRL1 paralogs largely overlap, they show unique properties regarding the speed and capacity of PSI oxidation during light stress. Our results suggest that PGRL1 duplication provided an evolutionary opportunity to diversify control of photoprotection and allow for dynamic range of protective responses to environmental stresses. Employing the two paralogs in a cell-specific manner in NADP-ME C_4_ photosynthesis conceivably contributed to the broader suite of adaptations enabling NADP-ME species to survive in high-stress environments.

## Results

### *PGRL1* paralogs diverged after genome duplication early in grasses evolution

To resolve the evolutionary history of PGRL1, we aligned *PGRL1* sequences from publicly available annotated genomes across the grass family, as well as selected monocot and dicot outgroups. The phylogenetic tree of *PGRL1* genes showed two distinct clades within the grass family which were designated as *PGRL1α* and *PGRL1β* (Fig. 1)*. PGRL1β* was absent in monocot outgroups *Musa acuminata* and *Joinvillea ascendens*, as well as in eudicot outgroups *Arabidopsis thaliana*, *Flaveria trinervia*, and *Artemisia annua* (Fig. 1). This indicates that functionally redundant *PGRL1Α* and *PGRL1Β* in *A. thaliana* both belong to the *PGRL1α* group. *Streptochaeta angustifolia,* which diverged from the main branch of grasses before the evolution of *Pharus latifolius* (Seetharam *et al*., 2021), also had only *PGRL1α* (Fig. S1). However, *PGRL1β* was present in Pharus, PACMAD, and BOP clades suggesting that *PGRL1β* emerged in the common ancestor of *P. latifolius* and the PACMAD and BOP clades, corresponding with a hypothesised whole-genome duplication event around 98 MYA (Ma *et al*., 2021).

**Fig. 1.**
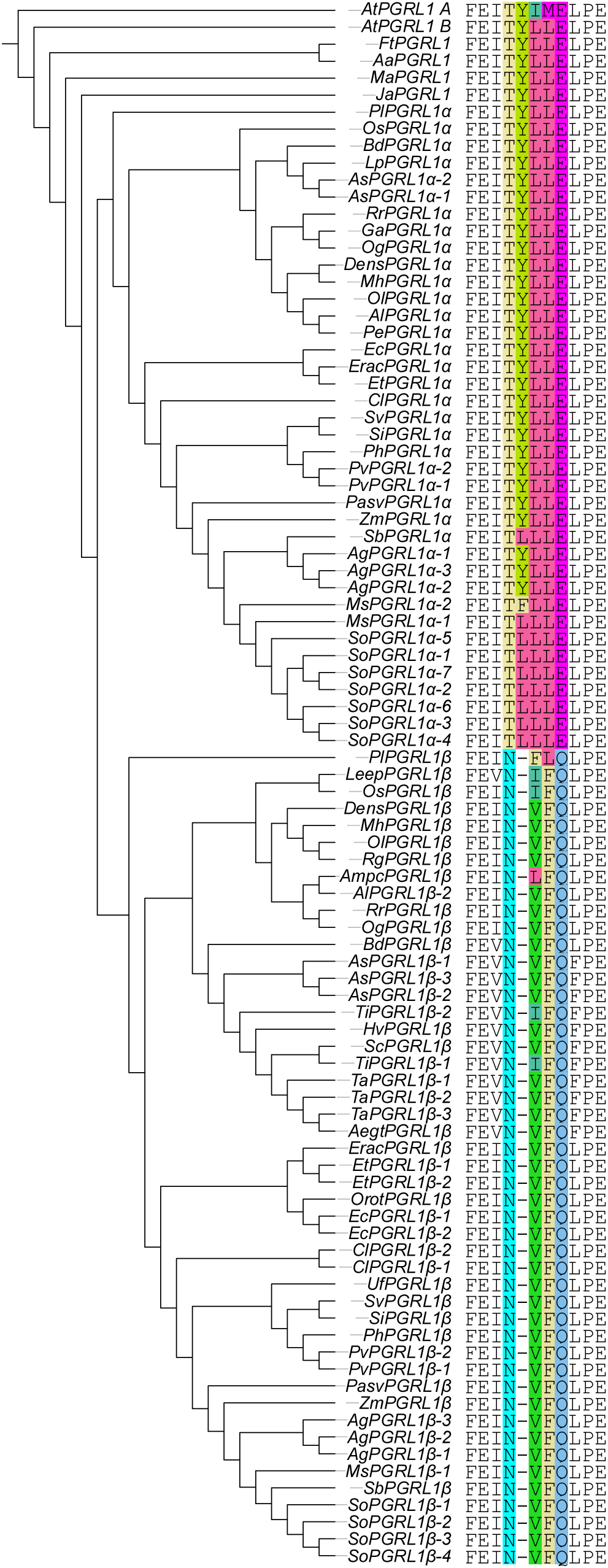
Sequence analysis of *PGRL1* homologs from grasses and other plant lineages. Phylogenetic tree of *PGRL1* generated through maximum-likelihood analysis based on the coding sequences of the mature proteins. The tree is rooted with *AtPGRL1Α*. In the alignment of the lumen regions [165-175 amino acids in *AtPGRL1Α*; DalCorso *et al*., 2008)], highlighted residues indicate sequence divergence among homologs. Additional copies of *PGRL1* within a species are suffixed as -1, -2, etc, which were assigned arbitrarily. *Aa, Artemisia annua; Aegt, Aegilops tauschii; Ag, Andropogon gerardii; Al, Ampelocalamus luodinaensis; Ampc, Ampelocalamus calcareus; As, Avena sativa; At, Arabidopsis thaliana; Ba, Bonia amplexicaulis; Bd, Brachypodium distachyon; Cl, Chasmanthium laxum; Dens, Dendrocalamus sinicus; Ec, Eleusine coracana; Erac, Eragrostis curvula; Et, Eragrostis tef; Ft, Flaveria trinerva; Ga, Guadua angustifolia; Hv, Hordeum vulgare; Ja, Joinvillea ascendens; Leep, Leersia perrieri; Lp, Lolium perenne; Ma, Musa acuminata; Mh, Melocanna humilis; Ms, Miscanthus sinensis; Og, Otatea glauca; Ol, Olyra latifolia; Os, Oryza sativa; Ot, Oropetium thomaeum; Pasv, Paspalum vaginatum; Pe, Phyllostachys edulis; Ph, Panicum hallii; Pl, Pharus latifolius; Pv, Panicum virgatum; Rg, Raddia guianensis; Rr, Rhipidocladum racemiflorum; Sb, Sorghum bicolor; Sc, Secale cereale; Si, Setaria italica; So, Saccharum officinarum; Sv, Setaria viridis; Ta, Triticum aestivum; Ti, Thinopyrum intermedium; Uf, Urochloa fusca; Zm, Zea mays.* Figure was generated using iTOL.

Among the 34 grass species examined, 24 species had at least one copy of both *PGRL1α* and *PGRL1β* (Fig. 1). In some species, multiple copies of *PGRL1α* or *PGRL1β* were observed, correlating with their genome architecture; for example, three copies of *PGRL1β* in *T. aestivum*, consistent with hexaploidy. However, no *PGRL1α* was found in 11 species and no *PGRL1β* in three species (Fig. 1, Table S1). Subsequent analysis suggested that *PGRL1α* in grasses was lost or rendered non-functional at least six times independently through random mutations in *Urochloa fusca*, *Oropetium thomaeum*, *Leersia perrieri*, *Raddia guianensis*, *Bonia amplexicaulis*, and Triticeae species (Fig. S2a). Potential *PGRL1β* loss-of-function mutations were observed in two woody bamboos (Fig. S3) and *Lolium perrene*, all of which lost functional transit peptides.

Next, we aligned the mature amino acid sequences (lacking chloroplast signal peptide) of PGRL1α and PGRL1β from 39 grass species (Fig. 1). Alignments yielded a pairwise positive BLSM62 score of 89%, with most variation due to single amino acid substitutions scattered across the sequence. However, a short region, previously shown to be exposed to the thylakoid lumen (DalCorso *et al*., 2008; Hertle *et al*., 2013), was consistently different between paralogs (Fig. 1). For PGRL1α this sequence was TYLLE, while for PGRL1β it was NVFQ. Further alignments of PGRL1α sequences from *S. viridis, A. thaliana*, *M. acuminata*, *F. trinerva*, *A. annua*, *Selaginella moellendorfii*, and *Chlamydomonas reinhardtii* showed that the TYLLE sequence was conserved as far back evolutionary as the lycophyte *S. moellendorffii*, with the green alga *C. reinhardtii* also having a similar sequence, TKLVE (Fig. S4). In contrast, the NVFQ sequence of PGRL1β was exclusive to grasses.

### *PGRL1α* and *PGRL1β* have distinct expression patterns

To gain insight into the functional differences between *PGRL1α* and *PGRL1β*, we performed a meta-analysis of available RNA-seq datasets of M and BS tissue-enriched fractions from *Oryza sativa* (rice, C_3_), *Panicum virgatum* (NAD-ME C_4_), *Z. mays*, *S. bicolor*, and *S. viridis* (NADP-ME C_4_) (Supplemental File 1; John *et al*., 2014; Döring *et al*., 2016; Rao *et al*., 2016; Denton *et al*., 2017; Chotewutmontri & Barkan, 2021; Hua *et al*., 2021). The log_2_ of the ratio between *PGRL1α* and *PGRL1β* transcript abundances (*PGRL1α*/*PGRL1β*) or corresponding protein abundances was derived to facilitate comparison between different datasets, with negative values indicating prevalent *PGRL1β* and positive values – *PGRL1α* (Fig. 2a). In *O. sativa* and *P. virgatum*, transcript abundance of *PGRL1β* was greater than *PGRL1α* in both M and BS cells (*O. sativa* M: -1.78, BS: -1.69; *P. virgatum* M: -3.06, BS: -2.80). Interestingly, in all studied NADP-ME grasses, of which *S. viridis* has independently evolved C_4_ photosynthesis (Sage *et al*., 2011), *PGRL1α* was preferentially expressed in M cells, while *PGRL1β* in BS cells (Fig. 2a). *S. bicolor* had the most cell-preferential expression (M: 3.39, BS: -6.30) while *S. viridis* had the least (M: 0.59, BS: -1.63). Transcript abundance of *PGR5* did not differ between M and BS cells in *S. viridis*, *Z. mays*, or *S. bicolor* (Supplemental File 1; John *et al*., 2014; Döring *et al*., 2016; Denton *et al*., 2017).

**Fig. 2.**
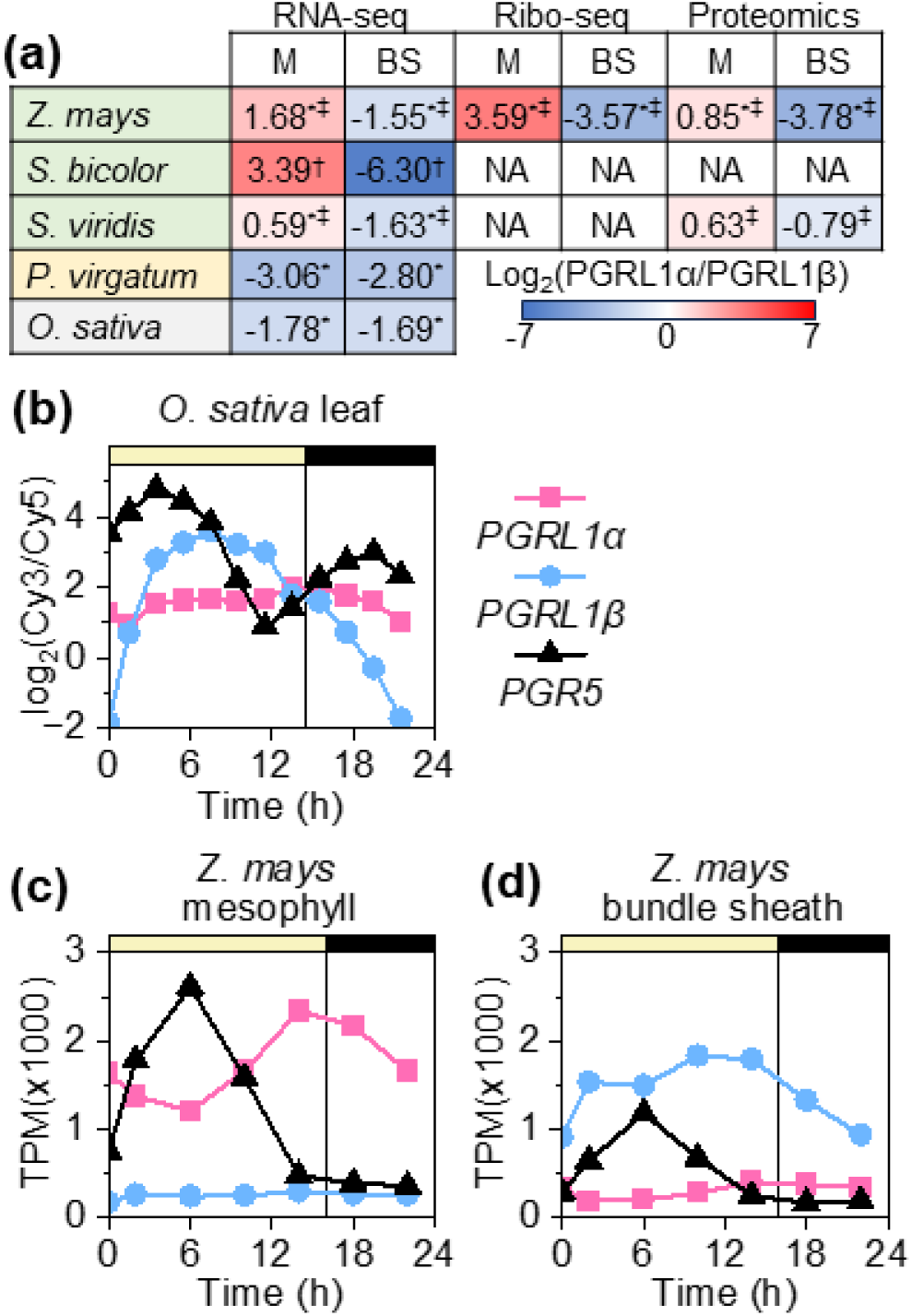
Expression analysis of *PGRL1α* and *PGRL1β* paralogs in grasses. (**a**) Tissue-specific levels of transcript abundance, actively-translated transcript abundance, and protein abundance in *Zea mays*, *Sorghum bicolor*, *Setaria viridis, Panicum virgatum,* and *Oryza sativa*. M, mesophyll BS, bundle sheath. Species are organised by the photosynthetic pathway: NADP-ME C_4_ (green), NAD-ME (yellow), or C_3_ (grey). Data shown as the log_2_ of the *PGRL1α*/*PGRL1β* ratio for transcript (or corresponding protein) abundances. Positive values indicate greater *PGRL1α* abundance (red) and negative values indicate greater *PGRL1β* abundance (blue). Transcriptional and translational data are sourced from RNA-seq and Ribo-seq datasets (John *et al*., 2014; Love *et al*., 2014; Döring *et al*., 2016; Rao *et al*., 2016; Denton *et al*., 2017; Chotewutmontri & Barkan, 2021; Hua *et al*., 2021), proteomics data for *Z. mays* are from Majeran *et al*. (2008); values for *S. viridis* were identified in this study. (**b**) Diurnal expression patterns of *PGRL1α*, *PGRL1β*, and *PGR5* in *O. sativa* from the microarray dataset (Sato *et al*., 2011; Sato *et al*., 2012). Expression levels are shown as log_2_ of the Cy3/Cy5 ratios with local sunrise set to 0 h. (**c, d**) Diurnal expression of *PGRL1α*, *PGRL1β*, and *PGR5* in M- and BS-enriched fractions of *Z. mays* obtained from the RNA-seq dataset of Borba *et al*. (2023). *PGR5* values shown for *Z. mays* are the sum of the two *PGR5* genes in *Z. mays* genome. TPM, transcripts per million. Light and dark phases are denoted by yellow and black bars. Symbols denote statistical significance from replicate-based tests: * log_2_(*PGRL1α*/*PGRL1β*) differed significantly from zero within a cell type; ‡ log_2_(*PGRL1α*/*PGRL1β*) was significantly different between M and BS cells; † pooled replicates in *S. bicolor* dataset precluding statistical testing (values shown descriptively). Full statistical details, source datasets, and replicate numbers used for this analysis are provided in the Supplemental File 1.

Analysis of cell-specific *Z. mays* Ribo-seq (Supplemental File 1; Chotewutmontri & Barkan, 2021) revealed cell-preferential translation of *PGRL1α* in M cells and *PGRL1β* in BS cells (M 3.59, BS -3.57). Furthermore, *Z. mays PGRL1β* transcript has been reported to contain a CACGCAC motif in the 5’-untranslated region (UTR), proposed to upregulate BS-specific translation (Chotewutmontri & Barkan, 2021). In contrast, *Z. mays PGRL1α* has lost a 5’-UTR and likely requires 5’-UTR-independent mechanisms of translational regulation (Majeran & van Wijk, 2009; Hardy & Balcerowicz, 2024). Analysis of cell-specific *Z. mays* proteomics data (Supplemental File 1; Majeran *et al*., 2008) showed PGRL1α preferentially in M cells (0.82) and PGRL1β preferentially in BS cells (-3.78). Our own mass spectrometry analysis of M and BS cell- enriched fractions from *S. viridis* found PGRL1α enriched in M cells (0.63) and PGRL1β enriched in BS cells (-0.79) (Fig. 2a; Supplemental File 1).

Analysis of publicly available *O. sativa* microarray data (Sato *et al*., 2011; Sato *et al*., 2012) revealed a strong diurnal expression pattern of *PGRL1β* but not *PGRL1α* (Fig. 2b). *PGRL1α* was the dominant transcript in the morning and evening while *PGRL1β* was the dominant transcript at midday. Diurnal expression patterns of *PGRL1α* and *PGRL1β* were also distinct in *Z. mays* (Borba et al., 2023). In M cells, *PGRL1α* expression was the lowest at 6 h of illumination and reached the peak at 14 h. In contrast, *PGRL1β* expression in BS cells was at its lowest during the dark period and peaked at 10 h of illumination. Expression of *PGRL1β* in M cells and *PGRL1α* in BS cells in *Z. mays* was consistently low, while PGR5 showed the same diurnal pattern in M and BS cells with a peak at 6 h, though was more abundant in M cells (Fig. 2c).

### PGRL1α is required for WT-level PSI oxidation at high light

To investigate specific roles of PGRL1 paralogs, we generated two null alleles for each gene of *S. viridis* (*pgrl1α-3*, *pgrl1α-10, pgrl1β-1,* and *pgrl1β-13*) using CRISPR-Cas9. Plants homozygous for each null allele (hereafter, *pgrl1α* and *pgrl1β* mutants) were selected by sequencing (Fig. S5), and the proteomic analysis confirmed the absence of the targeted paralog in each mutant (Fig. 3a,b). The two null alleles for each paralog exhibited no significant differences. Tissue-enriched proteomic analysis revealed WT-level abundance of the remaining PGRL1 paralog in each mutant, resulting in the total PGRL1 abundance in the mutants approximately half relative to WT but similar between each other (Fig. 3a,b). PGR5 abundance did not differ significantly between *pgrl1α* and *pgrl1β*, but was significantly lower in both mutants compared to WT.

**Fig. 3.**
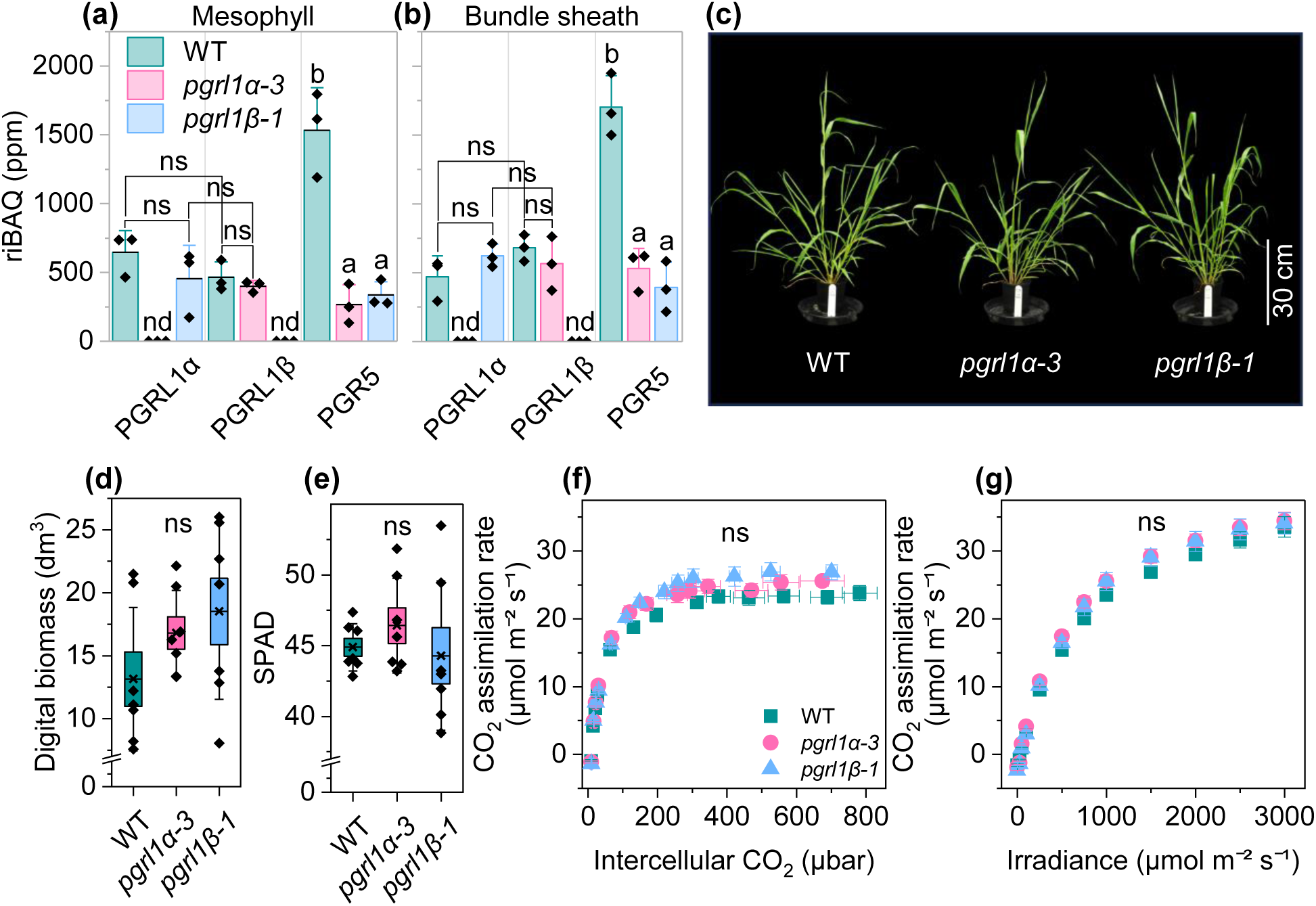
Growth and photosynthesis of *Setaria viridis* wild type (WT), *pgrl1α* and *pgrl1β* mutants grown under constant daylight (300 µmol m^−2^ s^−1^). (**a, b**) Relative intensity-based absolute quantification (riBAQ) values for protein abundances of PGRL1 paralogs in the mesophyll- or bundle sheath-enriched fractions. Mean ± SE, *n* = 3 biological replicates. Letters indicate significant differences between groups (one-way ANOVA with Tukey’s post hoc test at *P* < 0.05); ns, not significant; nd, not detected. (**c-e**) Growth phenotype, biomass, and relative chlorophyll content (SPAD) of four-week-old plants. Mean ± SE, *n* = 7 biological replicates. Letters indicate significant differences between groups (one-way ANOVA with Tukey’s post hoc test at *P* < 0.05); ns, not significant. (**f-g**) Response of the net CO_2_ assimilation rate to the intercellular CO_2_ partial pressure and irradiance. Mean ± SE, *n* = 5 biological replicates. Letters indicate significant differences between groups (two-way repeated measures ANOVA with Tukey’s post hoc test at P < 0.05); ns, not significant.

Growth, photosynthesis and relative chlorophyll content of the mutants grown in optimal conditions were similar to WT (Fig. 3c-e). No differences in CO₂ assimilation rates were found between genotypes across varying CO_2_ partial pressures or irradiances (Fig. 4f,g), with fitting of the gas-exchange curves showed no differences in biochemistry of photosynthesis (Table S2). *pmf* was similar between genotypes across irradiances (Fig. 4a). Analysis of chlorophyll fluorescence showed similar yields of photochemical and non-photochemical reactions within PSII at different irradiances across genotypes (Fig. S6). P700 (reaction centre of PSI) redox spectroscopy analysis showed that the quantum yield of photochemical reactions in PSI (φ_I_) was similar between all genotypes at all irradiances (Fig. 4b). However, *pgrl1α* had a lower quantum yield of non-photochemical reactions within PSI due to donor side limitation (φ_ND_), *i.e.* more reduced PSI, above 800 μmol photons m^−2^ s^−1^, compared to WT (Fig. 4c). At irradiances above 1,300 μmol photons m^−2^ s^−1^, *pgrl1α* also had a significantly higher quantum yield of non-photochemical reactions within PSI due to acceptor side limitation (φ_NA_) than WT (Fig. 4d).

**Fig. 4.**
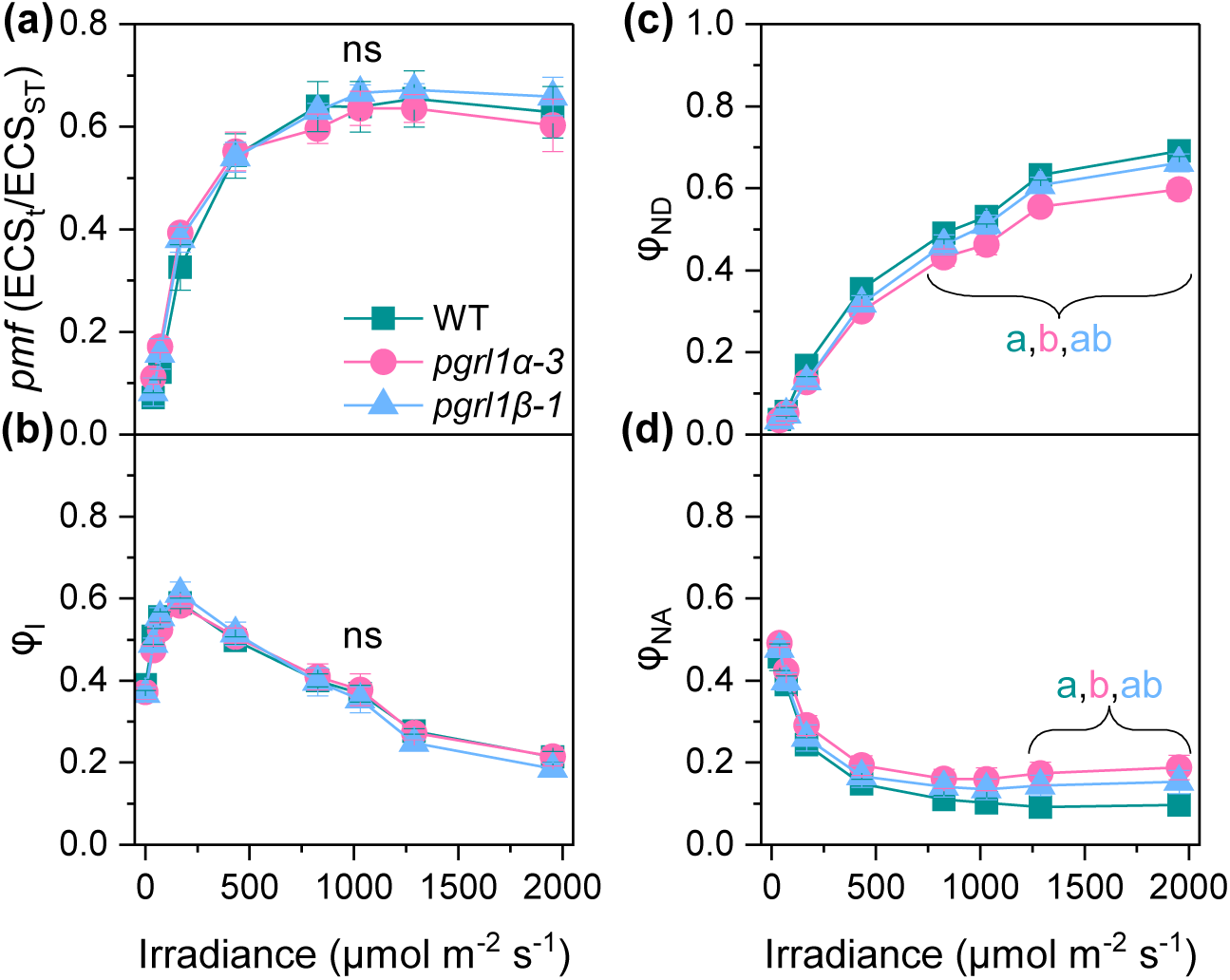
Light-response curves of *Setaria viridis* wild type (WT), *pgrl1α* and *pgrl1β* mutants grown under constant daylight (300 µmol m^−2^ s^−1^). (**a**) Proton motive force (*pmf*). (**b**) The photochemical yield of Photosystem I (PSI) (φ_I_). (**c**) The non-photochemical yield of PSI due to donor side limitation (φ_ND_). (**d**) The non-photochemical yield of PSI due to acceptor side limitation (φ_NA_). Mean ± SE, *n* = 5 biological replicates for (a), 8 for (b–d); letters indicate significant differences between groups (two-way repeated measures ANOVA with Tukey’s post hoc test at *P* < 0.05); ns, not significant.

### PGRL1β facilitates faster PSI oxidation

Next, we monitored leaf fluorescence and P700 redox state upon the shift of dark-adapted plants to 1,500 μmol photons m^−2^ s^−1^, i.e. during the induction of photosynthesis (Fig. 5). The maximum quantum efficiency of PSII (F_V_/F_M_) did not differ between genotypes, and the yields of photochemical and non-photochemical reactions within PSII were similar between the genotypes at 10 s and 60 s of illumination (Fig. 5a-c). The maximum oxidisable P700 (P_M_) was also similar between the genotypes but *pgrl1β* had higher φ_NA_ and lower φ_ND_ than both WT and *pgrl1α* at 10 s of illumination (Fig. 5d,e). After 60 s of illumination, PSI parameters were similar between all genotypes (Fig. 5f). Plotting the leaf P700^+^ signals normalised for P_M_ showed slower P700 oxidation in *pgrl1β* (Fig. 5g), and Boltzmann fitting of the curves revealed that P700 oxidation in *pgrl1β* was delayed for about 5 s compared to WT and *pgrl1α*, based on the higher midpoint parameter (*x₀*) (Fig. 5h). The slope factor (*dx*) did not differ between genotypes indicating that the rate of P700 oxidation was similar between genotypes (Fig. 5i).

**Fig. 5.**
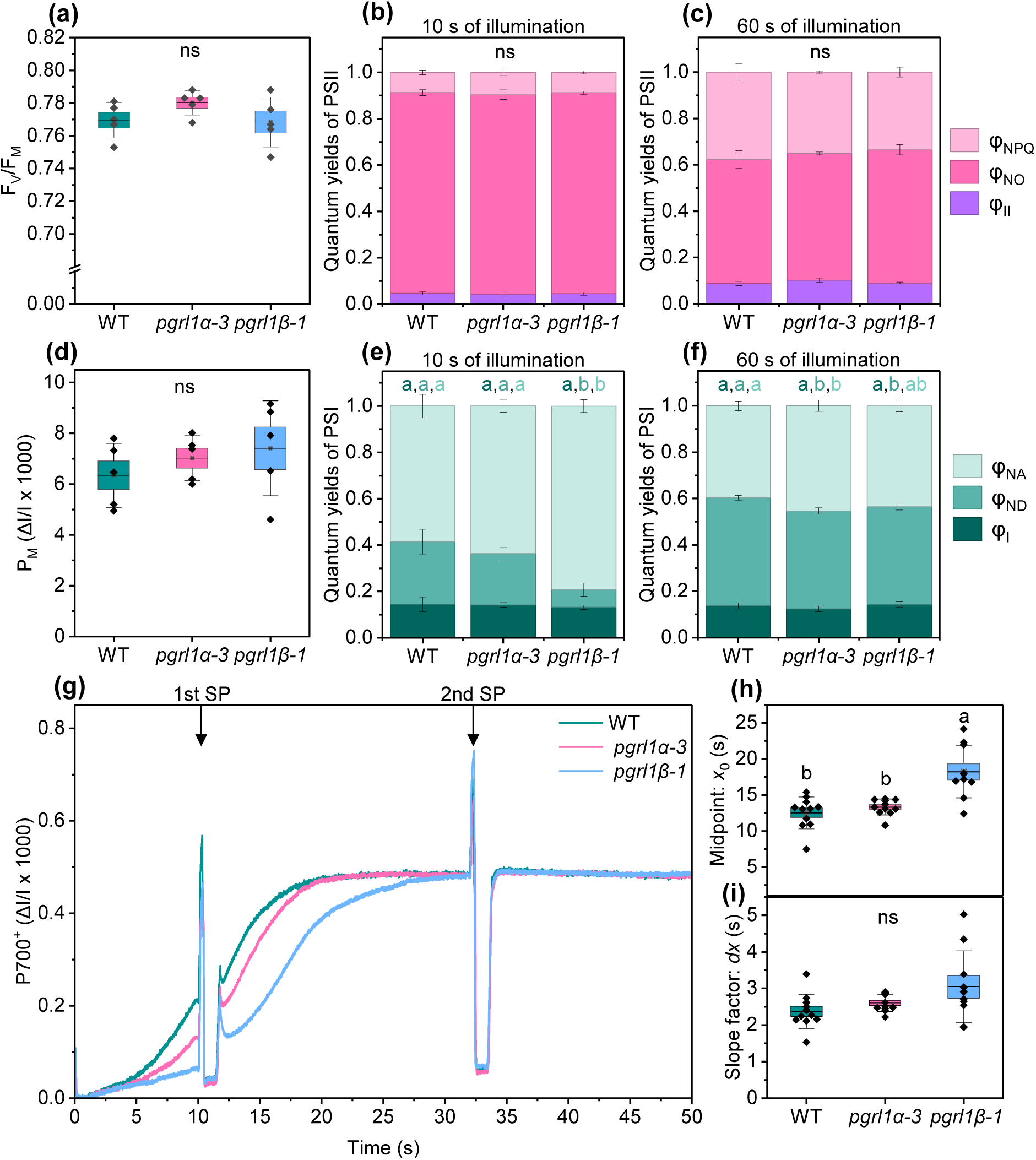
Induction of photosynthesis in *Setaria viridis* wild type (WT), *pgrl1α*, and *pgrl1β* mutants following a shift from dark to high light (1,500 µmol m^−2^ s^−1^). (**a**) Maximum Photosystem II (PSII) efficiency (F_V_/F_M_). (**b, c**) Partitioning of absorbed light energy within PSII between the effective quantum yield (φ_II_), the yield of non-regulated non-photochemical reactions (φ_NO_), and the yield of non-photochemical quenching (φ_NPQ_) at 10 and 60 s of the induction. (**d**) P_M_, the maximal photo-oxidisable P700, the reaction centre of Photosystem I (PSI). (**e, f**) Partitioning of absorbed light energy within PSI between the photochemical reactions (φ_I_), the non-photochemical reactions due to donor side limitation (φ_ND_) and the non-photochemical reactions due to acceptor side limitation (φ_NA_) at 10 and 60 s of the induction. (**g**) P700^+^ absorbance traces normalised by P_M_; arrows indicate saturating pulses (SP) applied; average of 10 biological replicates. (**h, i**) The midpoint (*x₀*) and the slope factor (*dx*) derived from Boltzmann fits of curves in (g). (a, d, h, i) Boxes show mean, SE, and SD, *n* = 5 biological replicates for (a, d), 10 for (h, i); letters indicate significant differences between groups (one-way ANOVA with Tukey’s post hoc test at *P* < 0.05); ns, not significant. (b, c, e, f) Mean ± SE, *n* = 8 biological replicates; letters indicate significant differences between groups (one-way ANOVA with Tukey’s post hoc test at *P* < 0.05); ns, not significant.

We also monitored the P700^+^ signal in leaves adapted to low light of 300 μmol photons m^−2^ s^−1^ and then shifted to 1,500 μmol photons m^−2^ s^−1^ (Fig. 6a). Plotting the P700^+^ signals normalised for P_M_ showed that, after 3 min at high irradiance, the P700^+^ level in *pgrl1α* was lower compared to WT and *pgrl1β*, which was supported by the lower φ_ND_ in *pgrl1α* at 180 s compared to WT and *pgrl1β* (Fig. 6a) and was in line with the lower φ_ND_ at high light seen in the light curve analysis (Fig. 4c). Normalising the curves for the P700^+^level at low irradiance (0) and after 45 s at high irradiance (1) to facilitate comparison of the kinetics showed that *pgrl1β* had slower oxidation of PSI compared to WT and *pgrl1α* (Fig. 6b). Furthermore, mutants had distinct levels of PSI damage under different types of light stress. After 1 h of constant high light (2,000 μmol m^−2^ s^−1^), *pgrl1α-3* showed significantly greater loss of P_M_ than both WT and *pgrl1β*, which did not differ from each other (Fig. 6c). In contrast, after 1 h of fluctuating light treatment (100 μmol photons m^−2^ s^−1^ for 115 s alternating with 2,000 μmol photons m^−2^ s^−1^ for 5 s), *pgrl1β* showed significantly greater loss of P_M_ than both WT and *pgrl1α*, which did not differ from each other (Fig. 6d).

**Fig. 6.**
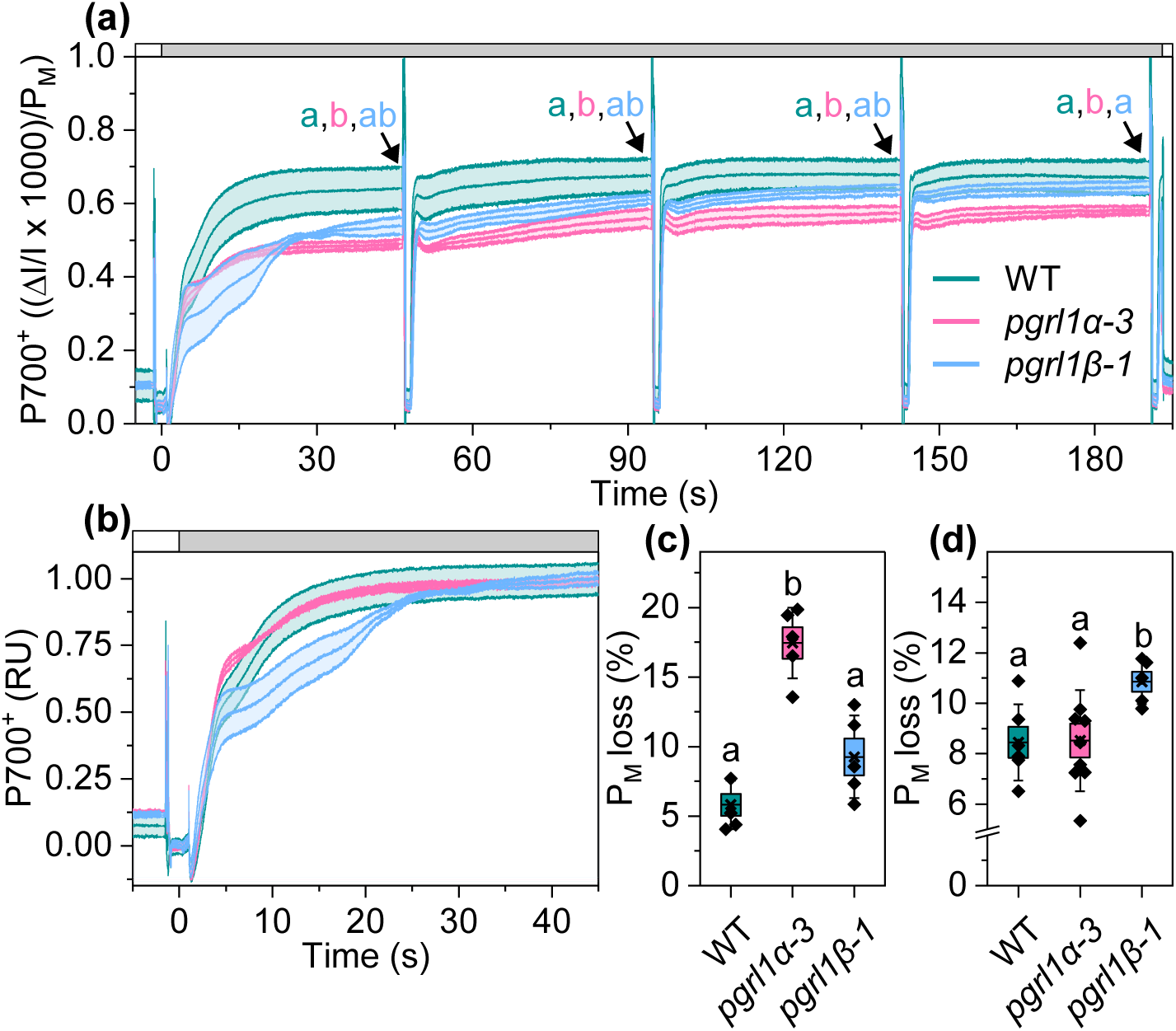
Responses of *Setaria viridis* wild type (WT), *pgrl1α* and *pgrl1β* mutants to light stress. (**a**) Oxidised reaction centre of Photosystem I (P700^+^) absorbance trace normalised for P_M_ (maximal photo-oxidisable P700) during a shift from low light (300 µmol m^−2^ s^−1^, white bar on top) to high light (1,500 µmol m^−2^ s^−1^, grey bar). Letters show significant differences between genotypes in φ_ND_, the yield of non-photochemical reactions in Photosystem I (PSI) due to donor side limitation, at each saturating pulse (two-way repeated measures ANOVA with Tukey’s post hoc test, *n* = 5 biological replicates, *P* < 0.05). (**b**) P700 oxidation kinetics from (a) normalised for P700⁺ levels at low light (0) and at 45 s of high light (1); RU, relative units. (**c**) Loss of P_M_ after 1 h of high light treatment (2,000 µmol m^−2^ s^−1^). (**d**) Loss of P_M_ after 1 h of fluctuating light treatment (100 µmol m^−2^ s^−1^ for 115 s alternating with 2,000 µmol m^−2^ s^−1^ for 5 s). (c, d) Boxplots show mean, SE and SD; letters indicate significant differences between groups (one-way Welch’s ANOVA with a Games-Howell post hoc test at *P* < 0.05); (c) *n* = 5; (d) *n* = 6 (WT), 9 (*pgrl1α*), 5 (*pgrl1β*).

### Protein models of PGRL1α and PGRL1β suggest structural basis for functional differences

To understand how PGRL1 paralogs provide distinct modes of photoprotection, we used AlphaFold3 to predict structures of the *S. viridis* PGRL1α and PGRL1β monomers (Fig. 7a). Consistent with prior experimental studies (DalCorso *et al*., 2008; Hertle *et al*., 2013), both paralogs had a transmembrane region and a large hydrophilic region on the stromal side. The Trxm4-sensitive cysteines responsible for PGRL1 dimerisation, and a rubredoxin-like domain involved in Fe^2+^ or Zn^2+^ binding, were structurally conserved between PGRL1α and PGRL1β. The key difference was found in the region exposed to lumen, where TYLLE in PGRL1α formed an alpha helix which was not observed in PGRL1β (Fig. 7a).

**Fig. 7.**
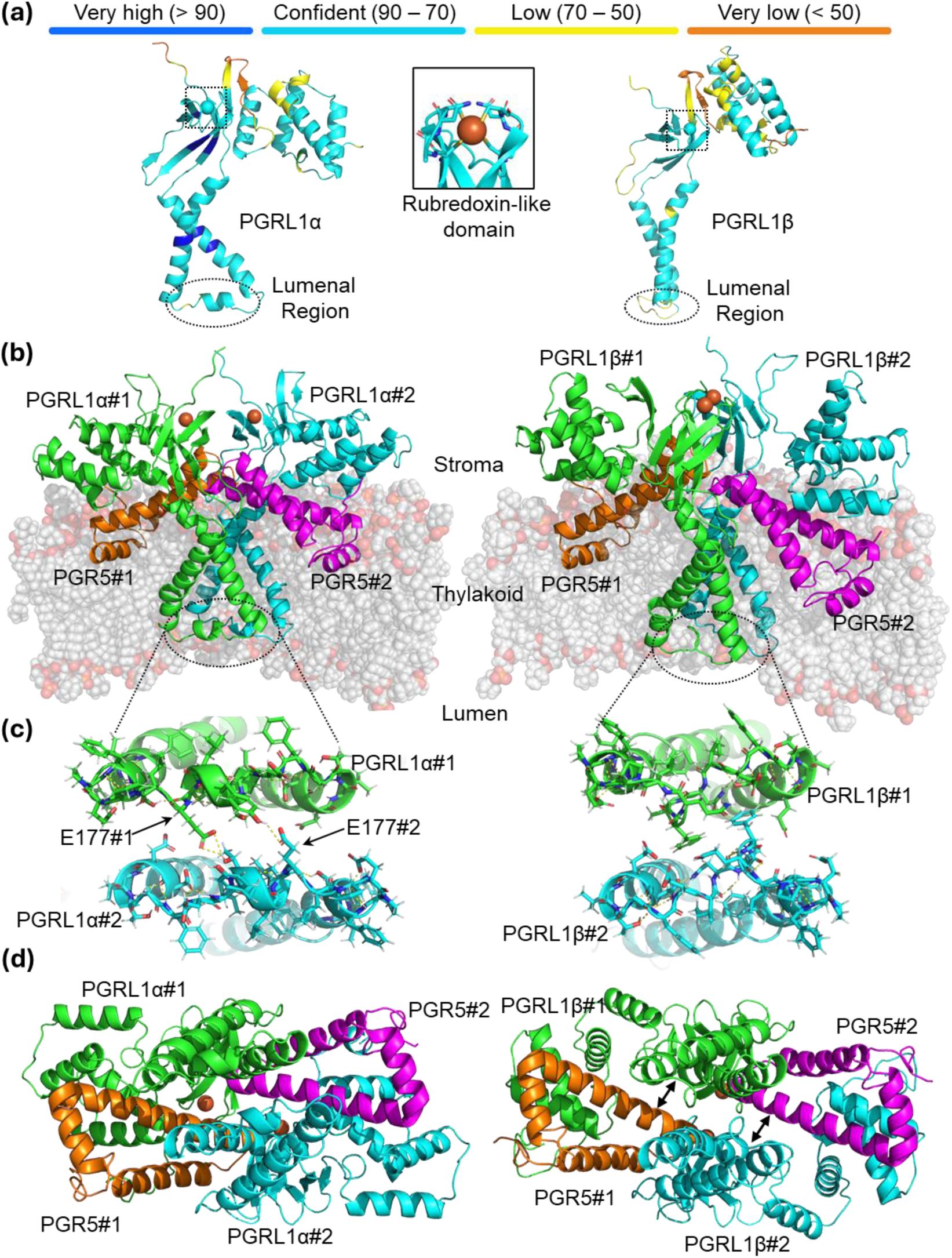
Structural models of PGRL1α and PGRL1β. (**a**) Predicted structures of PGRL1α and PGRL1β generated by AlphaFold3, with the rubredoxin-like domain in the inset. Predicted local distance difference test (pLDDT) confidence levels for each residue are indicated by colour, representing per-atom confidence on a scale from 0 to 100. (**b**) Membrane association models of PGR5-PGRL1α heterodimer-dimer (left) and PGR5-PGRL1β heterodimer-dimer (right). (**c**) Lumen and transverse view of the interface between PGRL1 monomers in PGR5-PGRL1α (left) and PGR5-PGRL1β (right) heterodimer-dimers. Hydrogen bonds are shown in yellow. E177 residues of each PGRL1α monomer are indicated. (**d**) Lumen view of PGR5-PGRL1α heterodimer-dimer (left) and PGR5-PGRL1β heterodimer-dimer (right) without the membrane. Arrows indicate a gap between the monomers.

To gain further insights, we generated AlphaFold3 models of a dimer of the PGR5-PGRL1 heterodimer complex, with either PGRL1α or PGRL1β, at neutral pH and with reduced cysteines, and assembled these complexes into a model membrane (Fig. 7b). PGRL1 is known to dimerise and interact with PGR5, providing rationale for modelling PGR5-PGRL1 heterodimer-dimers *in silico* (DalCorso *et al*., 2008; Rühle *et al*., 2021; Chaturvedi *et al*., 2024). Heterodimer-dimers formed a T-shaped structure, with each PGR5 binding in a hydrophobic pocket beneath the rubredoxin-like domain of one PGRL1 monomer, covered by the alpha helix ‘arm’ of the other PGRL1 monomer. This configuration was similar for both paralogs (Fig. 7b). The distances between the rubredoxin-like domains of the two PGRL1 monomers, however, were markedly different between paralogs. In PGRL1α, the domains were positioned side by side, 17.1 angstroms apart, whereas in PGRL1β, the domains faced each other with a shorter distance of 10.7 angstroms.

Both heterodimer-dimer models showed interactions between the lumen regions of PGRL1 monomers (Fig. 7c). In PGRL1α, hydrogen bonds were predicted between E177 of monomer #1 and T173 of monomer #2, and between E177 of monomer #1 and Y-174 of monomer #2. In PGRL1β, no hydrogen bonds were observed, but F173 of the NVFQ regions formed hydrophobic interactions between monomers. A transverse view showed that the PGRL1α and PGRL1β heterodimer-dimers had a different packing arrangement (Fig. 7d), and that the key differing regions were accessible from the thylakoid lumen (Fig. S7).

## Discussion

C_4_ photosynthesis has independently evolved over 60 times (Sage *et al*., 2011). Because C_4_ confers substantial advantages over conventional C_3_ photosynthesis in hot and dry conditions, there is strong interest in the evolutionary processes that led to its emergence (Sun *et al*., 2026). Different C_4_ subtypes achieve similarly high assimilation rates despite notable biochemical differences, highlighting the flexibility of C_4_ metabolism (Furbank, 2011; Fan *et al*., 2025). However, NADP-ME is the most efficient subtype under optimal conditions (Ehleringer & Pearcy, 1983; Arce Cubas *et al*., 2023) and is also the most studied. Consequently, NADP-ME is a common target for efforts to engineer C_4_ photosynthesis into C_3_ crops (Ermakova *et al*., 2021a). Achieving this goal requires a deeper understanding of the functional rationale and molecular mechanisms underlying cell-differentiation of photosynthetic processes. Interestingly, all three NADP-ME grasses studied here, representing two independent origins of C_4_ photosynthesis, have convergently evolved cell-specific expression of *PGRL1* paralogs (Fig. 2), strongly suggesting this arrangement is advantageous for C_4_ function.

To test whether the cell-specific arrangement of PGRL1α and PGRL1β is beneficial for C_4_ photosynthesis, we generated *S. viridis* mutants lacking either paralog. Cell-level proteomics confirmed that the *pgrl1α* mutants expressed only PGRL1β in both M and BS cells, while the *pgrl1β* mutants expressed only PGRL1α in both cell types. Despite the mutants having half the total PGRL1 and significantly lower PGR5 abundance compared to WT (Fig. 3a,b), *pmf* and φ_NPQ_ in both *pgrl1α* and *pgrl1β* were similar to WT levels (Fig. 4a, Fig. S6), confirming that either paralog was competent in regulating PGR5 and establishing low lumen pH to facilitate qE.

Despite similar levels of qE and PGR5 between mutants, PGRL1 paralogs each provided a distinct speed and capacity of PSI oxidation. Lower φ_ND_, the fraction of PSI in the oxidised state, in *pgrl1α* under steady-state high irradiance (Fig. 4c) and a greater loss in P_M_ after 1 h of high light treatment (Fig. 6c) indicates that PGRL1α in mesophyll cells was necessary for keeping PSI sufficiently oxidised at continuous high light. In contrast, lower φ_ND_ at 10 s of illumination and a delay in P700 oxidation in *pgrl1β* during the induction of photosynthesis (Fig. 5e,g), as well as slower P700 oxidation kinetics during the shift from low to high light (Fig. 6b) and a greater loss in P_M_ after a 1 h of fluctuating light (Fig. 6d), showed that PGRL1β was required in BS cells for prompt PSI oxidation. Although the relationship between φ_ND_ and photosynthetic control is complex (Degen & Johnson, 2024), based on the strong link between PGR5-PGRL1 and photosynthetic control, we suggest that observed variations in φ_ND_ in the mutants could be attributed to differences in the regulation of photosynthetic control. A more intricate tuning of photosynthetic control than by lumen pH alone has been proposed, with regulation also thought to involve stromal redox state (Degen & Johnson, 2024). This is seen in tobacco plants with Ferredoxin:NADP^+^ oxidoreductase knock- down, which show uncoupling of qE (and by extension lumen pH) and photosynthetic control (Hald *et al*., 2008). The similar decoupling seen here in PGRL1 mutants indicates that PGRL1 paralogs may differentially modulate photosynthetic control beyond just lowering lumen pH.

Differences in cell-specific electron transport provide clues into why distinct modes of PSI photoprotection conferred by PGRL1 paralogs are required in M and BS cells of NADP-ME grasses. M cells have a high abundance and activity of PSII and therefore predominantly operate linear electron flow (LEF), while BS cells have lower PSII activity and instead operate high CEF through the chloroplast NADH dehydrogenase-like complex (Ermakova *et al*., 2021b; Ermakova *et al*., 2024). In CEF-dominant systems, while the overall rate of electron transport can be high, it is conceivable that the electron flux on the donor side does not substantially exceed the electron flux at the acceptor sides of PSI, meaning that a limitation in electron acceptors is unlikely. Comparatively, in LEF-dominant systems, the capacity of the PSI acceptor side is largely governed by the activity of stromal metabolic reactions, which may be limited, for example, during increases in irradiance due to the time required for enzyme activation (Wang *et al*., 2021; Arrivault *et al*., 2025). Therefore, in M cells, if a sudden increase in irradiance stimulates PSII activity, electron influx on the donor side of PSI can significantly exceed the capacity of the acceptor side. Notably, however, this electron influx likely occurs more slowly in LEF than in CEF, as electrons must move from PSII, which resides in stacked grana, to PSI residing in the lamellae (Wójtowicz *et al*., 2025). Consequently, fast but low capacity photoprotection conferred by PGRL1β is advantageous in CEF-dominant BS cells, whereas high-capacity protection conferred by PGRL1α is preferable in M cells to account for high electron flux from PSII, even at the expense of lower speed. Interestingly, the ratio of *PGRL1β* vs *PGRL1α* transcript abundance in BS cells also correlated well with the relative PSII abundance between species (Fig. 2): it was highest in sorghum with lowest PSII content and lowest in *S. viridis* with highest PSII content (Meierhoff & Westhoff, 1993; Ermakova *et al*., 2021b), further supporting that the ratio is tuned to the photosystems’ composition.

The lumen-exposed sequences of PGRL1α and PGRL1β may underlie the distinct modes of photoprotection conferred by the paralogs. Our protein models suggested that the lumen regions of PGRL1α monomers were linked via hydrogen bonds formed by the negatively charged E177 (Fig. 7c), potentially stabilising the dimer. If this is the case, activation of PGR5 could require not only reduction of the stromal cysteines by Trxm4 but also lumen acidification to protonate the glutamates. This putative pH-sensing role is reminiscent of the function of lumen-exposed glutamates in PsbS that facilitate structural changes under low pH to trigger qE (Liguori *et al*., 2019; Krishnan-Schmieden *et al*., 2021). In contrast, lumen regions of PGRL1β lack the corresponding glutamate and appear to be linked by hydrophobic interactions (Fig. 7c). During increases in irradiance, *pmf* is initially dominated by the electric potential before equilibrating toward a proton gradient (Davis *et al*., 2017; Wilson *et al*., 2021), suggesting that responses of reactions depending on lumen pH might be slightly delayed. Therefore, lumen pH-independent PGRL1β might enable faster photoprotection under light transients compared to PGRL1α.

Having both PGRL1 paralogs enabling distinct modes of photosynthetic control is evolutionary advantageous. Our phylogenetic analysis suggests a duplication event after the origin of grasses gave rise to PGRL1β. While whole-genome duplications are common in plants, non-essential duplicates are often lost during subsequent speciation or through random mutations (Rieseberg & Willis, 2007). The persistence of both PGRL1α and PGRL1β in most grasses for up to ∼98 million years therefore implies a selective benefit of the dual system. Consistent with this, analysis of *S. viridis* mutants indicates that the paralogs confer complementary functions and together provide both fast and high-capacity photoprotection at the leaf level. Cooperative systems with complementary functions in regard to speed/capacity or affinity/flux are common in biological systems and provide broader operational range; a typical example is membrane transporters (Levy *et al*., 2011). Notably, however, some grasses have lost one paralog, including the group that contains wheat, rye, and barley. It will be important to determine how the absence of PGRL1α in these species shapes the regulation of photosynthetic control relative to C_3_ grasses that retain both paralogs.

## Conclusion

The duplication of PGRL1 in grasses enabled functional divergence, with PGRL1β evolving as a rapid-response regulator of photoprotection, while PGRL1α retained its ancestral role in slower but high-capacity protection. These differences in function likely arose from a divergence in the lumen region, with PGRL1α activity dependent on pH regulation of a conserved glutamate, which is absent in PGRL1β. The co-option of PGRL1 paralogs into M and BS cells by NADP-ME C_4_ grasses supports distinct modes of photoprotection offered by the PGRL1 paralogs and suggests that it’s a requirement for efficient NADP- ME C_4_ photosynthesis. As modern agriculture relies on open-field environments, often with fluctuating irradiance, introducing PGRL1β into non-grass crops such as canola, potato and soybean might offer enhanced resilience to light stress, presenting a new potential avenue for crop improvement.

## Materials & Methods

### Phylogenetic analysis

All phylogenetic analyses were performed in Geneious Prime (Geneious, 2024). *PGRL1* sequences were obtained from the Phytozome (Goodstein *et al*., 2011), Gramene (Tello-Ruiz *et al*., 2022), and CoGe (Lyons & Freeling, 2008) databases. Grass species were selected based on availability of a complete genome assembly and an RNA-seq based transcriptome annotation. *PGRL1* genes were identified using a predicted-protein search. For the phylogenetic tree, coding sequences (CDS) corresponding to the mature protein were utilised. First, peptide sequences were processed using TargetP 2.0 (Almagro Armenteros *et al*., 2019) to predict chloroplast transit peptides. Mature protein regions were then mapped back to their corresponding CDS. Gene IDs can be found in Table S1.

Protein alignment was conducted using MUSCLE 5.1 with default settings (Edgar, 2022). Multiple sequence alignment of CDS was carried out using the MAFFT v7.490 algorithm (Katoh *et al*., 2002; Katoh & Standley, 2013) with the BLOSUM62 scoring matrix, a gap open penalty of 1.53, and an offset value of 0.123. The phylogenetic tree was generated using RAxML v8.2.11 (Stamatakis, 2014) based on the MAFFT alignment, with the following parameters: Nucleotide Model: GTR CAT, Algorithm: Rapid Bootstrapping and search for best-scoring ML tree, Number of starting trees: 100, Parsimony random seed: 1. The resulting RaxML bootstrapped tree was finalised and annotated in iTOL (Letunic & Bork, 2021).

### *PGRL1* expression analyses

Expression of *PGRL1s* and corresponding proteins in *Zea mays*, *Sorghum bicolor*, *Setaria viridis, Panicum virgatum* and *Oryza sativa* was examined by investigating publicly available RNA-seq (John *et al*., 2014; Döring *et al*., 2016; Rao *et al*., 2016; Denton *et al*., 2017; Hua *et al*., 2021; Borba *et al*., 2023), Ribo-seq (Chotewutmontri & Barkan, 2021), and proteomics (Majeran *et al*., 2008) datasets. Data were manually extracted from supplementary files, and for each sample the log_2_ of *PGRL1α*/*PGRL1β* transcript (or protein) abundance ratio was calculated to allow comparison across datasets. Full data sources, per-replicate values, and detailed statistical methods are provided in Supplemental File 1. Detailed methods on tissue enrichment, RNA extraction and sequencing, and data processing are provided in original publications.

### Plant growth conditions

*S. viridis* plants were germinated on rooting medium as described in Osborn *et al*. (2016), then transferred to soil in 0.5 L pots with seed-raising mix and 5 g L^−1^ of slow-release Osmocote fertilizer (Scotts, Australia). Plants were grown in a chamber with ambient CO₂, temperatures of 28°C (day) and 22°C (night), and 60% humidity, and a constant light regime (380 µmol m^−2^ s^−1^, 16 h photoperiod). All measurements and analyses were performed on the youngest fully expanded leaves of three-to-four-week-old plants.

### Mass spectrometry analyses

To obtain tissue-specific enriched protein fractions from WT *S*. *viridis*, M-enriched proteins were isolated via the leaf rolling method (Furbank *et al*., 1985), and BS strands were obtained by differential grinding (Furbank & Badger, 1983), followed by homogenisation with mortar and pestle to extract proteins as described in Ermakova *et al*. (2019). Protein extracts were digested with trypsin following the filter-aided sample preparation protocol (Wiśniewski *et al*., 2009). Briefly, 10 μg of protein in each sample was suspended in lysis buffer [50 mM tris(hydroxymethyl)aminomethane (Tris), 25 mM NaCl, 20 mM Ethylenediaminetetraacetic acid (EDTA), 1% sodium dodecyl sulphate (SDS), 5 mM dithiothreitol (DTT)], heated to 95°C for 5 minutes, sonicated, and centrifuged at 16,000 g for 5 minutes. Lysate was loaded onto a Amicon Ultra-0.5 Centrifugal Filter (Millipore) with 20 μL of urea buffer (8 M urea, 50 mM ammonium bicarbonate) to retain the total protein. Vacuum was applied to facilitate buffer elution from the columns. Following an initial wash, 15 μL of chloroacetamide was added to the urea buffer, and three additional washes with urea buffer (without chloroacetamide) were performed. Proteins were digested on the filter with 120 μL of 50 mM ammonium bicarbonate and 5 μL of 400 ng μL^−1^ trypsin for 8 h at 37°C. Digested peptides were collected with 50 μL of water. Peptides were then dried using a speed vac, resuspended in 0.1% formic acid, and sonicated. Samples were loaded onto an Orbitrap Fusion Mass Spectrometer (Thermo Fisher Scientific, Waltham, Massachusetts, USA). Data were analysed using MaxQuant software (Cox & Mann, 2008), and relative protein abundances were calculated using an intensity-based absolute quantification (iBAQ) approached based on Schwanhäusser *et al*. (2011), modified to use the number of observed peptides, rather than theoretically observable peptides. The observed iBAQ values were then converted to relative iBAQ (riBAQ) values by dividing each protein’s iBAQ value by the total iBAQ sum in that replicate, such that riBAQ values represent each protein’s fractional contribution to the total protein abundance in the sample. Minimal contamination of tissue-enriched fractions was identified using malic enzyme and phosphoenolpyruvate carboxylase as signature proteins for BS and MS cells respectively.

### CRISPR-Cas9 mutant generation

*S. viridis* cv. ME034V-1 plants with null *pgrl1α* and *pgrl1β* alleles were created using CRISPR/Cas9 gene-editing as described in Ermakova *et al*. (2024). The genomic and coding sequences of *S. viridis PGRL1α* and *PGRL1β* were obtained from Phytozome and can be found in Supplemental Table 1. Cas9 was targeted to the first exons of each gene using gRNAs selected with CRISPOR (Concordet & Haeussler, 2018) and assembled into a synthetic polycistronic gene according to Xie *et al*. (2015). The Golden Gate cloning system (Engler *et al*., 2014) was used to assemble the construct which was transformed into *S. viridis* using *Agrobacterium tumefaciens* strain AGL1 (Osborn *et al*., 2016). Editing of the target genes was confirmed by sequencing in T_1_ plants and the progenies of plants lacking T-DNA with homozygous null alleles were used in all experiments. Two independent null alleles for both *pgrl1α* and *pgrl1β* were obtained and verified by sequencing and proteomics.

### Gas-exchange analysis

Gas-exchange analysis was performed using the portable gas-exchange system LI-6800 (LICOR, Lincoln, Nebraska USA) equipped with a 6800-01A fluorometer head. For all measurements 90% red/10% blue actinic light was used with 25°C leaf temperature, 55% relative chamber humidity, flow rate of 500 µmol s^−1^, flow pressure of 0.1 kPa and fan speed of 10,000 RPM. For CO_2_ response curves, leaves were clamped into the chamber and equilibrated at 400 μmol mol^−1^ CO_2_ on the reference side and 1,000 μmol m^−2^ s^−1^ irradiance for 15 min. CO_2_ assimilation rates were then recorded at CO_2_ partial pressures from 0 to 1,600 μmol mol^−1^ at 3-min intervals. For light response curves, leaves were equilibrated at 400 μmol mol^−1^ CO_2_ and 1,000 μmol m^−2^ s^−1^ irradiance for 15 min. CO_2_ assimilation was then recorded with irradiance progressively increasing from 0 to 3000 μmol m^−2^ s^−1^ at 3-min intervals. CO_2_ and light response curves were fitted using the biochemical model of C_4_ photosynthesis parameterised for *S. viridis* (von Caemmerer, 2021; Woodford *et al*., 2026).

### Leaf spectroscopic and fluorescence analyses

To measure light responses of PSI and PSII quantum yields, simultaneous measurements of chlorophyll fluorescence and P700 redox spectroscopy were performed using a DUAL-PAM/F (Heinz Walz, Effeltrich, Germany) under 635 nm red actinic light with red light saturating pulses (300 ms, 12,000 μmol m^−2^ s^−1^). P700 redox state was assessed by detecting absorbance of the P700^+^ at 830 nm with a dual wavelength unit (830/875 nm). Leaves were dark-adapted for 30 min before applying a saturating pulse to determine the maximum (F_M_) and minimum (F_0_) levels of fluorescence. The maximum quantum yield of PSII (F_V_/F_M_) was calculated as (F_M_ - F_0_) / F_M_. Following this, the maximal P700^+^ signal (P_M_) was recorded by applying a saturating pulse at the end of an 8-s far-red light (720 nm) illumination, with the minimal P700^+^ signal (P_0_) recorded directly following the saturating pulse. After determining dark-adapted parameters, leaves were light-acclimated to 300 µmol m^−2^ s^−1^ for 15 min.

Light response curves were recorded by progressively increasing the actinic light from 0 to 2,000 µmol m^−2^ s^−1^ and applying a saturating pulse at the end of every 120-s illumination period. Upon the application of each saturating pulse, F_M_’ (the maximum level of fluorescence under light) and F (the fluorescence level before the application of a saturating pulse) were recorded. This enabled the monitoring of the partitioning of absorbed light energy within PSII between the photochemical [φ_II_ = (F_M_’ – F) / F_M_’] and non-photochemical reactions, including the regulated [φ_NPQ_ = (F_M_ – F ^’^) / F_M_] and non-regulated (φ_NO_ = F / F_M_) fractions (Kramer *et al*., 2004). The photochemical yield of PSI [φ_I_ = (P_M_’ – P) / (P_M_ – P_0_)], the non-photochemical yield of PSI due to acceptor side limitation [φ_NA_ = (P_M_ – P_M_’) / (P_M_ – P_0_)] and the non-photochemical yield of PSI due to donor side limitation [φ_ND_ = (P – P_0_) / (P_M_ – P_0_)] were calculated as described in Klughammer and Schreiber (2008).

PSI and PSII quantum yields during the photosynthetic induction were monitored simultaneously using a Dual KLAS-NIR (Heinz Walz, Effeltrich, Germany) under a 635 nm red actinic light with saturating pulses of red light (300 ms, 12,000 μmol m^−2^ s^−1^). Leaves were dark-adapted for 30 min before determining F_V_/F_M_ and P_M_. After that, leaves were shifted to high light (1,500 µmol m^−2^ s^−1^) for 2 min, with saturating pulses applied at 10, 30, 60, 90, and 120 s and quantum yields of PSI and PSII calculated as described earlier. To analyse P700 oxidation kinetics, P700^+^ absorbance traces were baseline corrected to 0 and normalised for P_M_ for each sample. The midpoint (*x₀*) and slope factor (*dx*) derived from Boltzmann fits of P700 oxidation kinetics were determined using OriginPro (OriginLab, Northampton, MA).

To monitor P700 oxidation during a low-to-high light transition, after determination of P_M_, leaves were acclimated to 300 µmol m^−2^ s^−1^ for 15 min before being shifted to high light (1,500 µmol m^−2^ s^−1^), with saturating pulses (300 ms, 12,000 µmol m^−2^ s^−1^) applied every 45 s. P700⁺ absorbance traces were baseline corrected to 0 and normalised for P_M_ for each sample. For comparison of the kinetics, traces were normalised to the minimum P700⁺ level at low light (0) and the maximum at 45 s of high light (1).

To assess PSI damage under light stress, P_M_ was determined before and after light stress treatments. For treatments, leaves were clamped into the LI-6800 (using 90% red/10% blue actinic light) at 25°C, 55% relative humidity, 400 µmol mol^−1^ CO₂. Leaves were exposed for 1 h to either a fluctuating light regime (100 µmol m^−2^ s^−1^ for 115 s alternating with 2,000 µmol m^−2^ s^−1^ for 5 s) or a constant high light of 2,000 µmol _m-2 s-1._

Electrochromic shift (ECS) signals were monitored with a Dual-PAM-100 equipped with the P515/535 emitter-detector module (Heinz Walz, Effeltrich, Germany). ECS was estimated from the absorbance changes between 550 and 520 nm and normalised for the amplitude of the ECS response to a saturating pulse (20 μs, 14,000 μmol m^−2^ s^−1^) measured from dark-adapted leaves (ECS_ST_). Changes in the ECS signal were measured during 3-min illumination periods of increasing irradiance. After each illumination period, the light was switched off for 5 s and the kinetics of ECS decay were analysed to determine the proton motive force (*pmf*). *Pmf* was estimated as the total change in amplitude of the ECS signal upon the light-to-dark transition (ECS_t_) normalised for ECS_ST_ (Takizawa *et al*., 2007).

### Plant phenotyping

Rapid phenotyping was performed using MultispeQ V2.0 (PhotosynQ, East Lansing, MI, USA) and the standard Photosynthesis RIDES 2.0 protocol which determines photosynthetic parameters at the incident irradiance (Kuhlgert *et al*., 2016). MultispeQ measurements were made under ambient conditions on plants grown under either a constant or fluctuating light environment. For fluctuating light, measurements were performed during the low irradiance periods. Digital plant biomass and leaf area was estimated using a PlantEye F600 Multispectral 3D scanner (Phenospex, Heerlen, Netherlands). Whole-plant imaging was performed on four-week-old plants to generate 3D point clouds, with digital biomass then calculated using voxel-based reconstructions.

### Protein modelling

Protein structures of monomers and complexes were built using the AlphaFold3 web server (Abramson *et al*., 2024). PGR5-PGRL1 heterodimer-dimer structures were modelled using the mature sequences for PGR5, PGRL1α and PGRL1β from *S. viridis* (Gene/Protein IDs found in Table S1) with a single Fe^2+^ ion per PGRL1 monomer and 50 palmitic acid molecules to provide a pseudo-membrane during structure prediction. Specific models were chosen for analysis from the AlphaFold3 output structures based on their pTM and ipTM scores. Model protein structures were embedded in a DOPC bilayer model as a proxy for a thylakoid membrane using the CHARMM-GUI server and membrane builder (Jo *et al*., 2008; Wu *et al*., 2014) and the OPM server (Lomize *et al*., 2011) to orient the protein with respect to the modelled membrane. Protein-membrane complexes were energy minimised in NAMD (Phillips *et al*., 2020) and inspected using PyMOL (Schrodinger, 2015).

## Acknowledgements

We thank Dr Adam Carrol and The Australian National University’s Research School of Biology/Research School of Chemistry Joint Mass Spectrometry Facility for protein mass spectrometry analysis. We thank Jacques Bouvier for advice and discussions. We acknowledge the use of the facilities, and scientific and technical assistance of the Australian Plant Phenomics Network, which is supported by the Australian Government’s National Collaborative Research Infrastructure Strategy (GA319462). We acknowledge support from the Australian Research Council (DP260100980, DP230100175, CE140100015, IC210100047).

## Authors declare no conflict of interest. Authors contribution

SJN, ME and RTF designed research. SJN and RW performed experiments. CJM generated membrane protein models. SJN and ME analysed data and wrote the manuscript. All authors discussed results and edited final manuscript.

