## Supplementary figures and tables for "Cell-preferential PGRL1 paralogs provide distinct modes of photoprotection in C_4_ photosynthesis"

**Supplementary materials**

**Table S1.** Accessions numbers of *PGRL1* genes studied in this work.

| **Acronym** | **Species** | **Gene** | **Gene ID** | **Gene** | **Gene ID** |
| --- | --- | --- | --- | --- | --- |
| Aa | *Artemisia annua* | *PGRL1* | Chromosome 7; 44,765-49,699 |  |  |
| Aegt | *Aegilops tauschii* |  |  | *PGRL1β* | AET5Gv21221900.2 |
| Ag | *Andropogon gerardii* | *PGRL1α-1* | AndgeH1.02BG087800.1 | *PGRL1β-1* | AndgeH1.01BG004300.1 |
|  |  | *PGRL1α-2* | AndgeH1.02AG108200.1 | *PGRL1β-2* | AndgeH1.01AG004100.1 |
|  |  | *PGRL1α-3* | AndgeH1.02CG092200.1 | *PGRL1β-3* | AndgeH1.01CG004700.1 |
| Al | *Ampelocalamus luodinaensis* | *PGRL1α* | Alu07Dg10900.2 | *PGRL1β-1* | Alu03Cg00420.1 |
|  |  |  |  | *PGRL1β-2* | Alu03Dg30180.2 |
| Ampc | *Ampelocalamus calcareus* | *PGRL1α* | Hca03Dg11500.3 | *PGRL1β* | Hca03Dg00380.1 |
| As | *Avena sativa* | *PGRL1α-1* | AVESA.00001b.r3.2Cg0001548.1 | *PGRL1β-1* | AVESA.00001b.r3.Ung0000396.1 |
|  |  | *PGRL1α-2* | AVESA.00001b.r3.2Dg0003409.1 | *PGRL1β-2* | AVESA.00001b.r3.7Cg0000037.1 |
|  |  |  |  | *PGRL1β-3* | AVESA.00001b.r3.4Dg0002573.3 |
| At | *Arabidopsis thaliana* | *PGRL1Α* | AT4G22890 |  |  |
|  |  | *PGRL1Β* | AT4G11960 |  |  |
| Ba | *Bonia Amplexicaulis* | *PGRL1α* | Bam07Ag07840.1 | *PGRL1β* | Bam03Ag19370.1 |
| Bd | *Brachypodium distachyon* | *PGRL1α* | Bradi1g52550.12 | *PGRL1β* | Bradi1g00940.1 |
| Cl | *Chasmanthium laxum* | *PGRL1α* | Chala.10G086100.1 | *PGRL1β-1* | Chala.03G290600.1 |
|  |  |  |  | *PGRL1β-2* | Chala.12G040300.1 |
| Cr | *Chlamydomonoas reinhardtii* | *PGRL1* | Cre07.g340200.t1.1 |  |  |
| Da | *Dioscorea alata* | *PGRL1* | Dioal.16G045900.1 |  |  |
| Dens | *Dendrocalamus sinicus* | *PGRL1α* | Dsi07Ag08820.1 | *PGRL1β* | Dsi03Ag00220.2 |
| Ec | *Eleusine coracana* | *PGRL1α* | ELECO.r07.7BG0597380.1 | *PGRL1β1* | ELECO.r07.3BG0252680.1 |
|  |  |  |  | *PGRL1β2* | ELECO.r07.3AG0207630.1 |
| Erac | *Eragrostis curvula* | *PGRL1α* | EJB05_12845 | *PGRL1β* | EJB05_03684 |
| Et | *Eragrostis tef* | *PGRL1α-1* | Et_5B_043645 | *PGRL1β-1* | Et_4B_038547 |
|  |  | *PGRL1α-2* | Et_5A_040970 | *PGRL1β-2* | Et_4A_034424 |
| Ft | *Flaveria trinerva* | *PGRL1* | Chromosome 7; 5,778,300 - 5,722,381 (reverse) |  |  |
| Ga | *Guadua angustifolia* | *PGRL1α* | Gan07Bg09810.2 | *PGRL1β* | Gan05Bg00110.1 |
| Hv | *Hordeum vulgare* |  |  | *PGRL1β* | HORVU5Hr1G124210.5 |
| Ja | *Joinvillea ascendens* | *PGRL1* | Joasc.10G046600 |  |  |
| Leep | *Leersia perrieri* | *PGRL1α* | LPERR08G08870 | *PGRL1β* | LPERR03G35530.1 |
| Lp | *Lolium perenne* | *PGRL1α* | KYUSg_chr2.31374 | *PGRL1β* | KYUSg_chr4.681 |
| Ma | *Musa acuminata* | *PGRL1* | GSMUA_Achr4T28830_001 |  |  |
| Mh | *Melocanna humilis* | *PGRL1α* | Mhu07Ag04200.1 | *PGRL1β* | Mhu03Bg00260.2 |
| Ms | *Miscanthus sinensis* | *PGRL1α-1* | MisinT443400 | *PGRL1β-1* | Misin16G024900 |
|  |  | *PGRL1α-2* | Misin04G108900 | *PGRL1β-2* | MisinT360900 |
| Og | *Otatea glauca* | *PGRL1α* | Ogl07Bg10220.1 | *PGRL1β* | Ogl03Cg23350.1 |
| Ol | *Olyra latifolia* | *PGRL1α* | Ol07g09230.1 | *PGRL1β* | Ol03g30000.1 |
| Ot | *Oropetium thomaeum* |  |  | *PGRL1β* | Oropetium_20150105_23595 |
| Os | *Oryza sativa* | *PGRL1α* | LOC_Os08g41460 | *PGRL1β* | LOC_Os03g64020 |
|  |  | *PGR5* | LOC_Os08g45190 |  |  |
| Pasv | *Paspalum vaginatum* | *PGRL1α* | Pavag02G081500 | *PGRL1β* | Pavag01G004100 |
| Pe | *Phyllostachys edulis* | *PGRL1α* | Ped07Dg06700.1 | *PGRL1β* | Ped03Cg00420.1 |
| Ph | *Panicum hallii* | *PGRL1α* | Pahal.2G103600 | *PGRL1β* | Pahal.9G004200 |
| Pl | *Pharus latifolius* | *PGRL1α* | Phala.03G097200 | *PGRL1β* | Phala.01G006700 |

| Pv | *Panicum virgatum* | *PGRL1α-1* | Pavir.2KG249300 | *PGRL1β-1* | Pavir.9KG015400 |
| --- | --- | --- | --- | --- | --- |
|  |  | *PGRL1α-2* | Pavir.2NG289400 | *PGRL1β-2* | Pavir.9NG016300 |
| Rg | *Raddia guianensis* | *PGRL1α* | Rgu07g15140.1 | *PGRL1β* | Rgu03g28870.3 |
| Rr | *Rhipidocladum racemiflorum* | *PGRL1α* | Rhi07Bg10380.1 | *PGRL1β* | Rhi03Cg00320.1 |
| Sa | *Streptochaeta angustifolia* | *PGRL1α* | Scaffold_1550;  66,500 – 69,500 |  |  |
| Sb | *Sorghum bicolor* | *PGRL1α* | Sobic.002G112400.6 | *PGRL1β* | Sobic.001G005200 |
| Sc | *Secale cereale* |  |  | *PGRL1β* | SECCE7Rv1G0454960 |
| Si | *Setaria italica* | *PGRL1α* | Seita.2G109800 | *PGRL1β* | Seita.9G004200 |
| Sm | *Selaginella moellendorfii* | *PGRL1* | 66389 |  |  |
| So | *Saccharum officinarum* | *PGRL1α-1* | SoffiXsponR570.05Ag111900.1 | *PGRL1β-1* | SoffiXsponR570.01Bg004900.1 |
|  |  | *PGRL1α-2* | SoffiXsponR570.5_9Ag316600.1 | *PGRL1β-2* | SoffiXsponR570.01Eg004400.1 |
|  |  | *PGRL1α-3* | SoffiXsponR570.05Bg100500.1 | *PGRL1β-3* | SoffiXsponR570.01Cg005200.1 |
|  |  | *PGRL1α-4* | SoffiXsponR570.05Bg101000.1 | *PGRL1β-4* | SoffiXsponR570.01Ag004400.1 |
|  |  | *PGRL1α-5* | SoffiXsponR570.05Cg105400.1 |  |  |
|  |  | *PGRL1α-6* | SoffiXsponR570.05Eg088400.1 |  |  |
|  |  | *PGRL1α-7* | SoffiXsponR570.05Gg110600.1 |  |  |
| Sv | *Setaria viridis* | *PGRL1α* | Sevir.2G115600 | *PGRL1β* | Sevir.9G004600 |
| Ta | *Triticum asetivum* |  |  | *PGRL1β-1* | Traes_4AL_A9E7CD7B8 |
|  |  |  |  | *PGRL1β-2* | Traes_5BL_283D01C08 |
|  |  |  |  | *PGRL1β-3* | Traes_5DL_89CF8AFCE |
| Ti | *Thinopryum intermedium* | *PGRL1α-1* | Thint.V1438200 | *PGRL1β-1* | Thint.14G0565500 |
|  |  | *PGRL1α-2* | Thint.14G0072400 | *PGRL1β-2* | Thint.14G0583200 |
|  |  | *PGRL1α-3* | Thint.V1087200 |  |  |
|  |  | *PGRL1α-4* | Thint.V1459800 |  |  |
| Uf | *Urochloa fusca* |  |  | *PGRL1β* | Urofu.9G004000.1 |
| Zm | *Zea mays* | *PGRL1α* | Zm00001d019454 | *PGRL1β* | Zm00001d034900 |
|  |  | *PGR5A* | Zm00001d031850 | *PGR5B* | Zm00001d049883 |

**Table S2.** Photosynthetic parameters of wild type (WT), *pgrl1α* and *pgrl1β* *S. viridis* plants fit from the CO_2_ response curves of CO_2_ assimilation (Fig. 3f). *V*_pmax_, phosphoenolpyruvate carboxylase activity; *V*_cmax_, Rubisco activity; *J*, electron transport rate required to sustain the measured CO_2_ assimilation rate [see details in Woodford *et al.* (2026)]. Mean ± SE, *n* = 5 biological replicates. No statistically significant differences were found between genotypes (one-way ANOVA with Tukey’s post hoc test at *P* < 0.05).

| **Parameter** | **WT** | ***pgrl1α-3*** | ***pgrl1β-1*** |
| --- | --- | --- | --- |
| ***J*** | 96.5 | 103.2 | 109.4 |
| (μmol m^-2^ s^-1^) | ±4.5 | ±3.5 | ±4.6 |
| ***V*_pmax_** | 62.0 | 70.6 | 60.1 |
| (μmol m^-2^ s^-1^) | ±5.5 | ±8.5 | ±3.2 |
| ***V*_cmax_** | 30.0 | 32.3 | 35.7 |
| (μmol m^-2^ s^-1^) | ±2.0 | ±2.5 | ±2.4 |

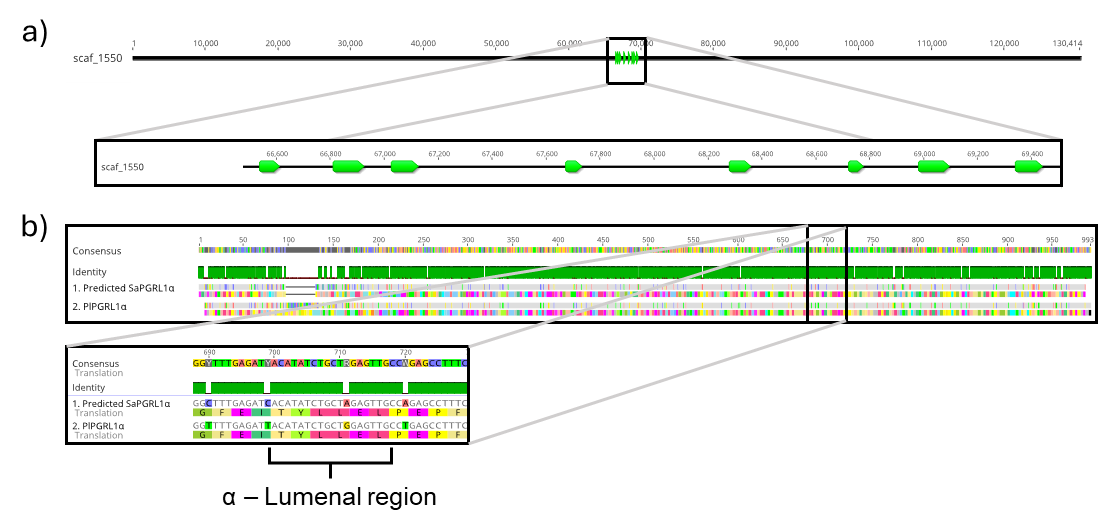

**Fig. S1.** Alignment of the predicted *PGRL1* gene from *Streptochaeta angustifolia* (Sa). **(a)** Alignment of *Pharus latifolius* (Pl) *PGRL1α* against the scaffold assembly of Sa. **(b)** MUSCLE alignment of the predicted Sa *PGRL1α* and Pl *PGRL1α*, highlighting the amino acid sequence of the lumenal region characteristic of PGRL1α.

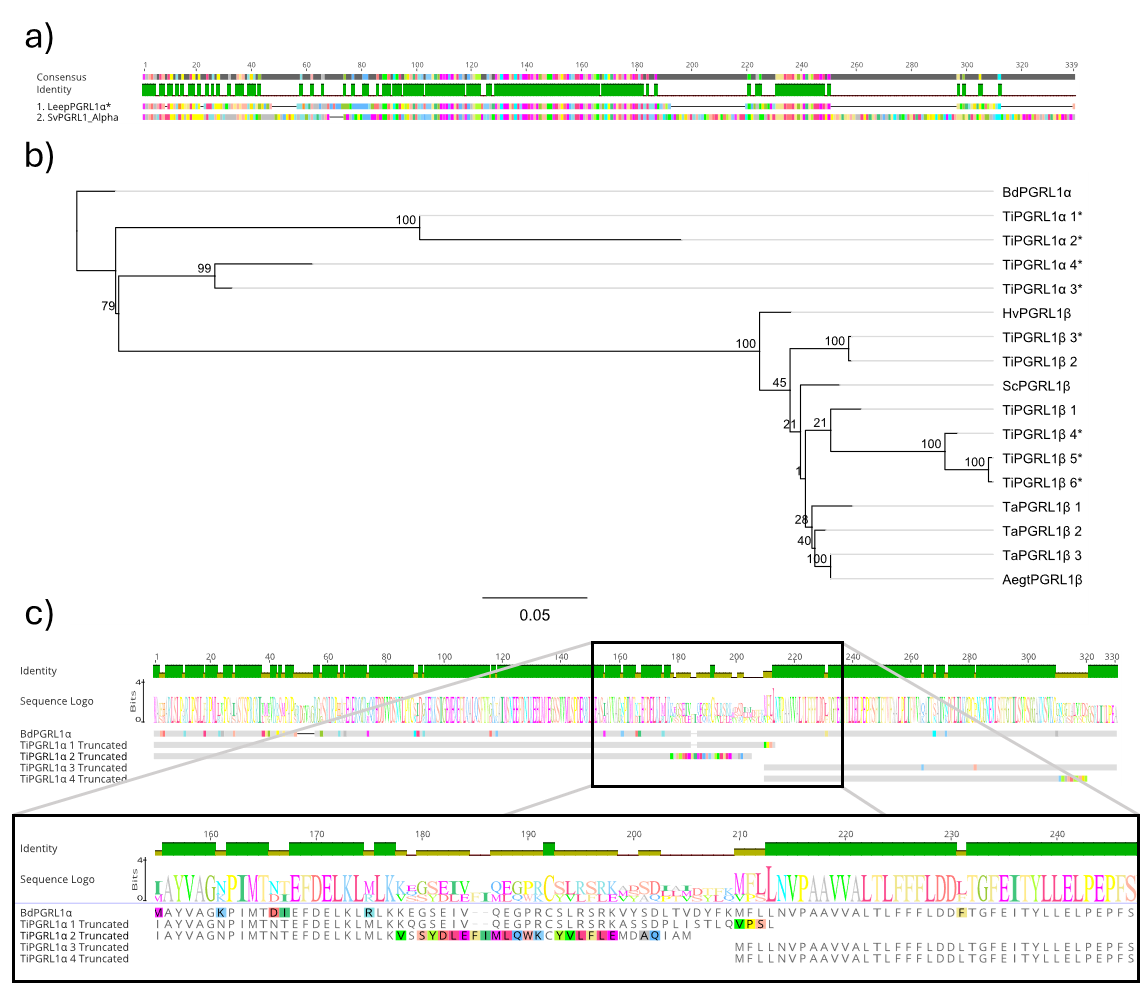

**Fig. S2.** Loss of functional *PGRL1α* genes across the Poaceae family. *PGRL1α* was lost or rendered non-functional at least six times independently in *Urochloa fusca*, *Oropetium thomaeum*, *Leersia perrieri*, *Raddia guianensis*, *Bonia amplexicaulis*, and all members of the Triticeae subfamily. In *U. fusca* (PEPCK C_4_ subtype) and *O. thomaeum* (NAD-ME C_4_ subtype), the absence of *PGRL1α* was determined by the lack of RNA-seq annotated sequences. **(a)** Alignment of *L. perrieri* (Leep) and *Setaria viridis* (Sv) PGRL1α amino acid sequences. The chloroplast transit peptide (mapped by TargetP 2.0) and multiple sections of the mature protein are missing in Leep PGRL1α. **(b)** Analysis of *PGRL1α* genes from the Triticeae: *Aegilops tauschii* (Aegt), *Hordeum vulgare* (Hv), *Secale cereale* (Sc), *Triticum aestivum* (Ta), and *Thinopyrum intermedium* (Ti). RAxML phylogenetic tree generated from MAFFT-aligned *PGRL1* genes within Triticeae, with *Brachypodium distachyon* (Bd) as an outgroup. Bootstrap values are shown at each node. Genes that do not encode a functional mature peptide are marked by asterisks. **(c)** Alignment of Ti PGRL1α showing incomplete protein sequences. The incomplete protein sequences were unlikely to be an artifact of poor RNA-seq annotation, as the genes belonged to different genomic regions: TiPGRL1α-2 (*Thint.14G0072400.1*) at Chr14:53647416..53649826 (reverse); TiPGRL1α-1 (*Thint.V1438200.1*) at ChrUN:119455789..119461794 (reverse); TiPGRL1α-3 (*Thint.V1087200.1*) at scaffold_698:425178..427956 (forward); TiPGRL1α-4 (*Thint.V1459800.1*) at ChrUN:137199120..137203790 (reverse).

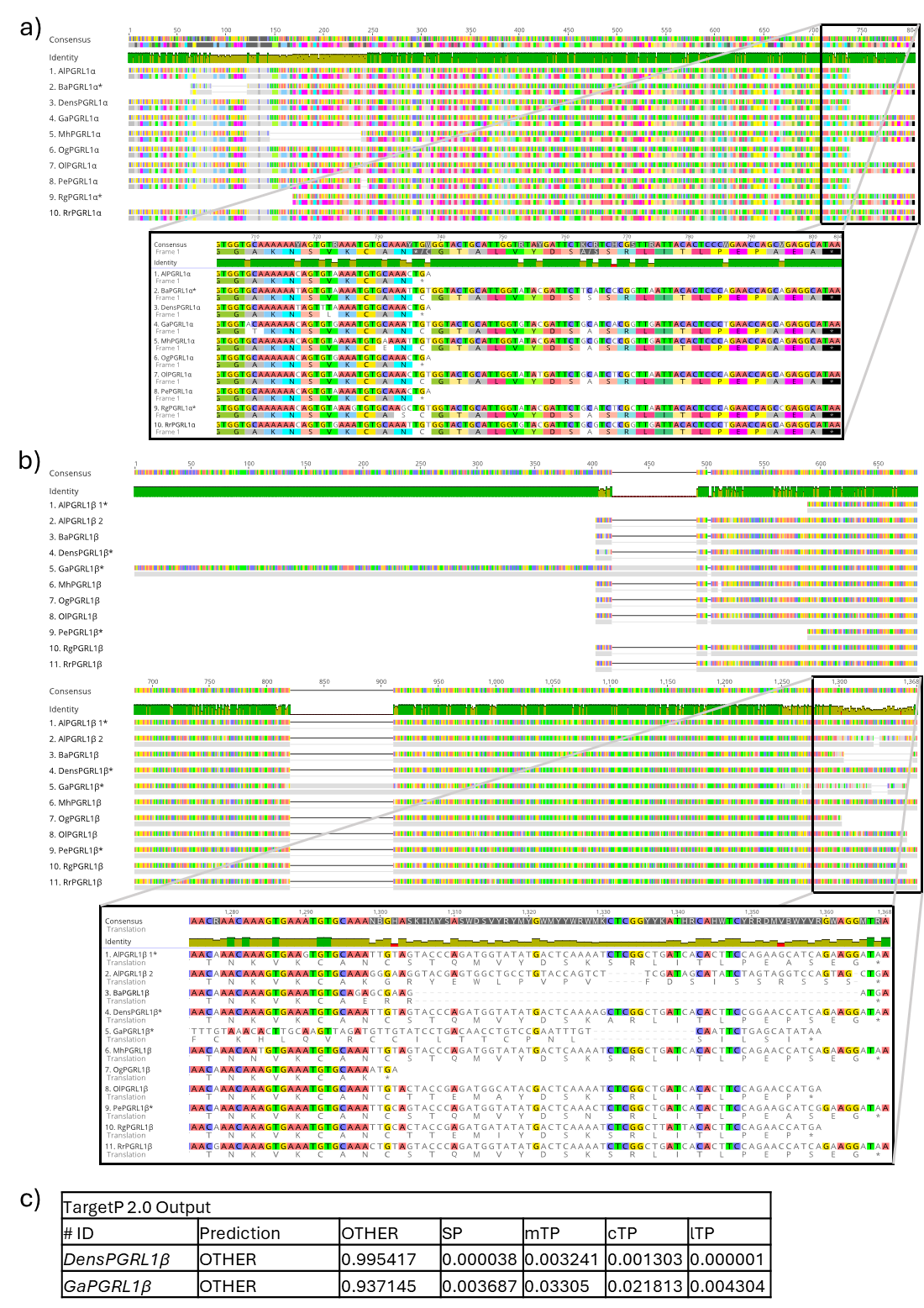

**Fig. S3.** Loss of functional *PGRL1* genes in the Bambusoideae subfamily. **(a)** MAFFT alignment of the *PGRL1α* genes and the corresponding amino acid sequences. Asterisks indicate genes that have lost functional chloroplast transit peptides. In *Melocanna humilis* (Mh) *PGRL1α*, a part of the coding sequence (CDS) in the beginning of the gene is missing, including the first cysteine involved in dimerisation. *Raddia guianensis* (Rg) and *Bonia amplexicaulis* (Ba) PGRL1α lack chloroplast transit peptides, suggesting that they are not localised to the chloroplast. *Ampelocalamus luodianensis* (Al), *Phyllostachys edulis* (Pe), *Dendrocalamus sinicus* (Dens), and *Otatea glauca* (Og) PGRL1α have premature stop codons and lack the final cysteine in the rubredoxin domain. According to Hertle et al. (2013), mutations at this cysteine do not affect iron binding in recombinant PGRL1 proteins, indicating that these proteins may retain some functional activity. **(b)** MAFFT alignment of the *PGRL1β* genes and the corresponding amino acid sequences. Asterisks indicate genes that have lost functional chloroplast transit peptides. Al and Pe PGRL1β lack chloroplast transit peptides due to later start codons, suggesting these proteins are not present in the chloroplast. Ba and Ga PGRL1β have premature stop codons and lack the final cysteine in the rubredoxin domain. **(c)** TargetP 2.0 outputs for Ga and Dens PGRL1β listed as probabilities, 1 being absolute certainty, 0 being no probability. TargetP 2.0 showed that both Ga and Rr *PGRL1β* lost a functional transit peptide, suggesting these proteins are not present in the chloroplast. OTHER – no signal peptide discovered, SP – signal peptide, mTP – mitochondrial transfer peptide, cTP – chloroplast transfer peptide, lTP – thylakoid lumen transfer peptide.

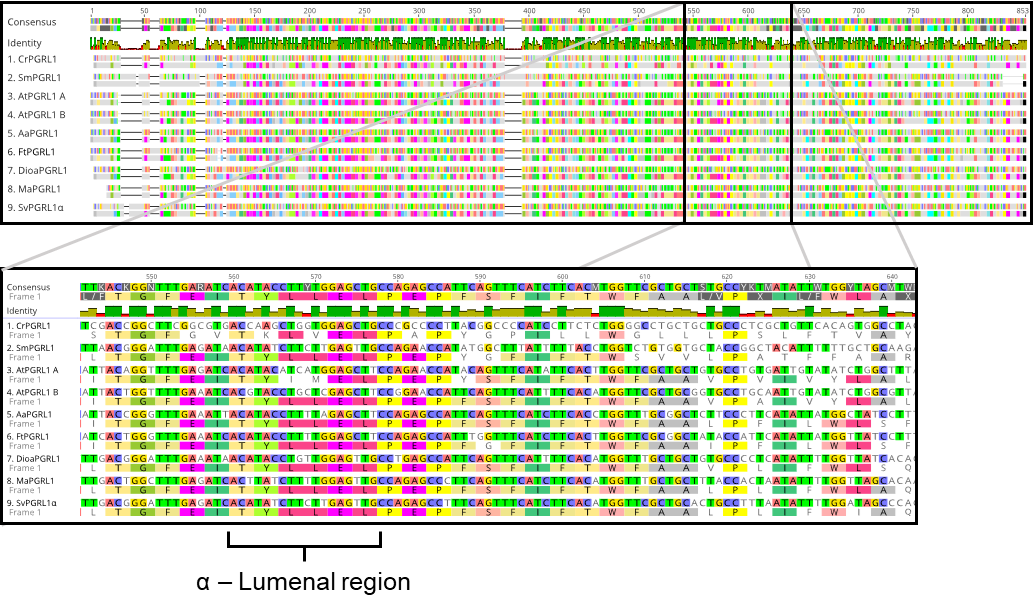

**Fig. S4.** MAFFT alignment of *PGRL1α* mature coding sequences and corresponding protein sequences from *Chlamydomonas reinhardtii* (Cr), *Selaginella moellendorfii* (Sm), *Arabidopsis thaliana* (At), *Artemisia annua* (Aa), *Flaveria trinerva* (Ft), *Dioscorea alata* (Dioa) and *Musa acuminata* (Ma) to *Setaria viridis* (Sv).

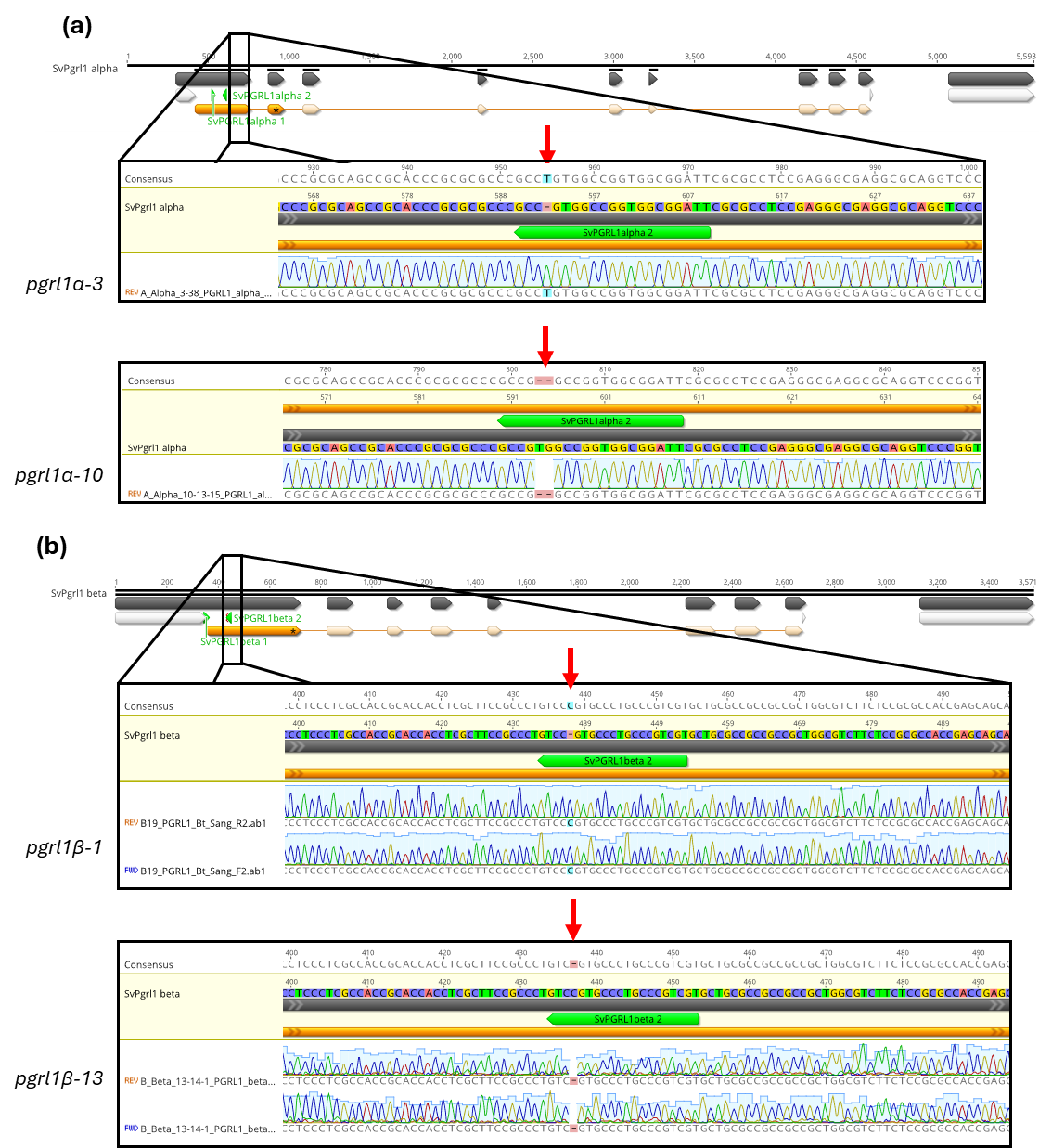

**Fig. S5.** Cas9-induced mutations in *pgrl1α-3, pgrl1α-10, pgrl1β-1* and *pgrl1β-13*. Gene models of *pgrl1α* (**a**) and *pgrl1β* (**b**) in *Setaria viridis* with coding sequence (CDS) annotated in orange. The introduced single nucleotide insertion, single deletion, or double deletion caused a frameshift which resulted in a premature stop codon. Sanger sequencing histograms confirm the introduced mutations.

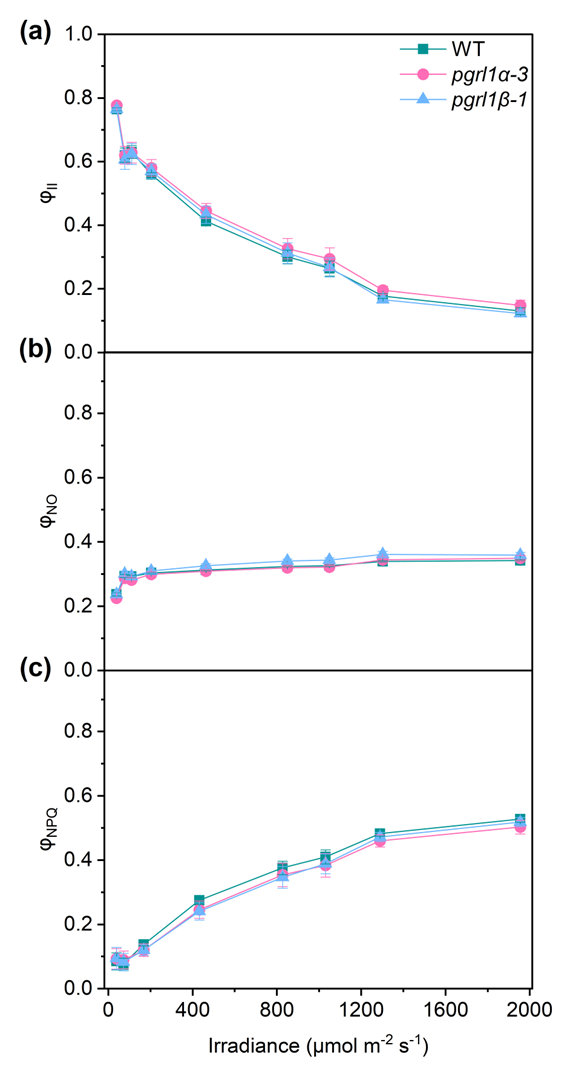

**Fig. S6.** Light-responses of (**a**) the effective quantum yield of PSII (φ_II_), (**b**) the yield of non-regulated non-photochemical energy dissipation (φ_NO_), (**c**) the yield of non-photochemical quenching (φ_NPQ_) in *Setaria viridis* wild type (WT), *pgrl1α* and *pgrl1β* mutant plants. Mean ± SE, *n* = 5 biological replicates; two-way repeated measures ANOVA found no statistical differences between genotypes.

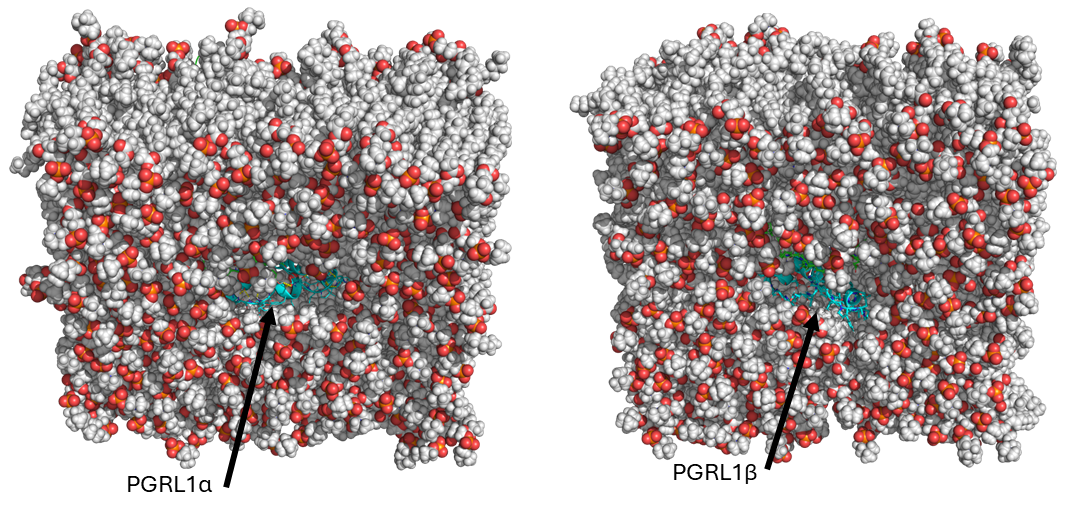

**Fig. S7.** View of PGRL1α-PGR5 and PGRL1β-PGR5 heterodimer-dimers modelled in the membrane from the lumen.
